# In situ cryo-ET of mammalian embryos reveals cytoplasmic lattices contain ubiquitin-charged E2-E3 ligase assemblies

**DOI:** 10.64898/2026.03.22.713481

**Authors:** Kashish Singh, Katarina Harasimov, Kathy K. Niakan, Andrew P. Carter

## Abstract

Cytoplasmic lattices (CPLs) are filamentous assemblies essential for mammalian embryonic development that regulate organelle organization, spindle assembly, and protein homeostasis. However, their molecular functions remain unclear. Here, we develop a strategy combining cryo-focused ion beam milling and cryo-electron tomography to resolve macromolecular complexes directly in mammalian embryos. Using this approach, we determine the in situ structure of CPLs within mouse embryos at ∼4.7 Å resolution. CPL filaments are built from multiple copies of fourteen proteins arranged into a ∼4.5 MDa repeating unit. These components form a scaffold that organizes a central cavity containing three complexes of the E2 ubiquitin-conjugating enzyme UBE2D, the E3 ligase UHRF1, NLRP14 and αβ-tubulin. We observe states in which UBE2D is conjugated to ubiquitin, along with structural rearrangements within the cavity consistent with regulated ubiquitin transfer. Together, our findings suggest that CPLs are large ubiquitin ligase assemblies and play a role in post-translational protein modification during early embryonic development.

## Introduction

Human embryonic development is a remarkably inefficient process. The majority of embryos perish before the onset of a clinically recognized pregnancy, primarily due to errors that arise during the first week after fertilization^1^. Early development is driven by maternally deposited factors, as the embryonic genome remains inactive for the first three days^2^. A critical set of maternal components are filamentous structures known as cytoplasmic lattices (CPLs)^3–6^. In mice, CPL loss results in developmental arrest shortly after fertilization^7–12^. In humans, mutations in CPL-associated proteins are linked to early embryonic arrest and infertility^13–15^, as well as miscarriage^14,16,17^ and multilocus imprinting syndromes^15,18–21^.

CPLs have been implicated in diverse cellular processes. Loss of CPL components disrupts organelle organization^22–24^ and spindle assembly during oocyte meiosis and early embryonic divisions^10,22,25,26^. CPL disruption has also been shown to impair proteostasis, leading to widespread changes in the abundance of diverse proteins, including ribosomal subunits, mitochondrial proteins, metabolic enzymes and cytoskeletal factors^27,28^. However, the molecular mechanisms through which CPLs affect these cellular processes remain unclear.

CPL filaments exhibit a highly regular ∼38 nm repeating unit and assemble into higher-order lattice-like arrays within the cytoplasm^27^. Genetic and light microscopy studies suggest that this repetitive structure is composed of the putative peptidylarginine deiminase PADI6 and the subcortical maternal complex (SCMC)^8,9,27^. The SCMC has been reported to contain an NLRP5-TLE6-OOEP complex together with multiple NLRP family proteins, including NLRP2, NLRP4B, NLRP4F and NLRP9, as well as KHDC3 and ZBED3 proteins^7,15,23,29–32^. In addition, more than 200 proteins whose abundance decreases upon CPL disruption have been proposed to be sequestered and stored within CPLs, where they are protected from degradation^27,28^. Interestingly, some of these proposed stored factors, such as NLRP14 and the E3 ubiquitin ligase UHRF1, are themselves required for CPL formation^11,12^. These observations raise key questions about which proteins are incorporated into CPL filaments, how they are organized within the repeating unit, and how this organization relates to CPL function.

Determining the molecular architecture of CPL filaments in their native environment has proven challenging. The scarcity of mammalian oocytes and embryos and the large size of these cells (∼80 µm in mice and ∼125 µm in humans) complicates in situ structural analysis by cryo-electron tomography, as preparing lamellae by cryo-focused ion beam milling must typically be performed manually, limiting throughput. Consequently, the only reported in situ reconstruction of CPLs, obtained from oocytes^27^, was limited to ∼30 Å resolution, insufficient for unambiguous protein identification.

Here, we develop a workflow enabling high-throughput cryo-focused ion beam milling and cryo-electron tomography for in situ structural analysis of early mammalian embryos. Using this approach, we determine the molecular architecture of cytoplasmic lattices within the native embryonic cytoplasm and show that they are built from a defined set of fourteen proteins. Their organization reveals that CPLs are assemblies that incorporate tubulin and ubiquitination machinery, providing a molecular framework for understanding their function during early development.

### In situ cryo-ET of early mouse embryos

We initially vitrified 6/8-cell stage mouse embryos directly on electron microscopy grids using a published cryopreservation protocol^27^ that utilizes 7.5% dimethyl sulfoxide and 7.5% ethylene glycol as cryoprotectants (Figure 1A and Figure S1A). We prepared lamellae from intact embryos by manual cryo-FIB milling and used cryo-ET to visualize macromolecular structures within the embryonic cytoplasm (Figure 1B and Figure S1B-C). We collected 36 tomograms from 14 embryos and reconstructed CPL filaments at 10 Å resolution by subtomogram averaging (Figure S1D). The resulting map closely resembled the previously reported 30 Å reconstruction from mouse oocytes^27^ (Figure S1E), indicating that CPL filament architecture is conserved between oocytes and cleavage-stage embryos.

**Figure 1.**
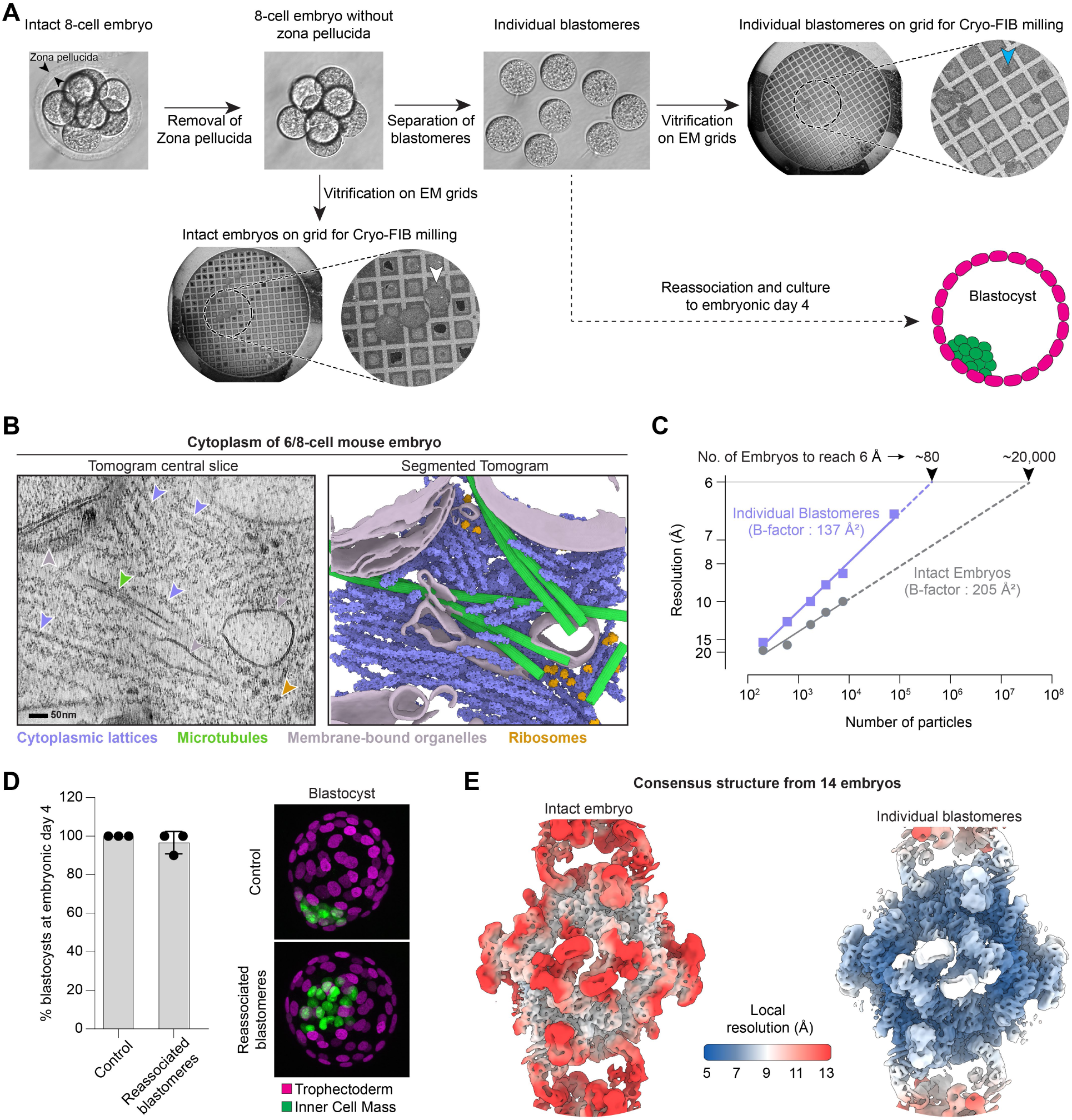
In situ structural investigation of CPLs from early mouse embryos. **(A)** Schematic for preparation of 6/8-cell stage mouse embryos for cryo-FIB milling directly or after separation into individual blastomeres. The white and green arrowhead highlights an example of an intact embryo and individual blastomere vitrified on an EM grid, respectively. **(B)** Central slice of a denoised tomogram depicting the cytoplasm within a blastomere of a 6/8-cell stage mouse embryos (left) with coloured arrowheads pointing to features in the corresponding segmentation of the tomogram (right). **(C)** Rosenthal-Henderson plot comparing datasets from intact embryos and individual blastomeres, demonstrating that data from individual blastomeres require fewer particles to achieve equivalent resolution. **(D)** Efficiency of blastocyst formation upon blastomere reassociation as shown in (A) compared to control samples where only the zona pellucida was removed (left). Immunofluorescence images of embryos at blastocyst stage from the two groups (right). Trophectoderm is marked by GATA3 (magenta), and inner cell mass by SOX2 (green). Independent replicates are shown as dots; mean ± s.d. are indicated. A minimum of five embryos were analysed per group in each replicate. **(E)** Subtomogram averaging density map of CPL repeat unit solved from intact embryos (left) and the corresponding map from individual blastomeres (right) coloured based on their local resolution.

However, the cryoprotectant concentrations required for vitrification of intact embryos reduced image contrast (Figure S2A), and the requirement for manual cryo-FIB milling limited the number of lamellae that could be prepared (Figure S2B). A Rosenthal-Henderson plot^33^ of resolution versus particle number shows that achieving ∼6 Å resolution, sufficient to resolve secondary structure features such as α-helices, would require an unfeasibly large amount of material (∼20,000 embryos) (Figure 1C).

To overcome these limitations, we developed a workflow in which embryos were gently separated into individual blastomeres immediately prior to vitrification (Figure 1A). Isolated blastomeres remained morphologically intact and formed blastocysts upon reassociation, showing that separation preserved cellular integrity and developmental potential (Figure 1A, 1D and S2C). Blastomere separation is routinely used for single-cell RNA sequencing^34^ and similar strategies have also recently enabled in situ structural analysis of cells isolated from complex tissues such as Drosophila ovaries^35^. Importantly, blastomere separation enabled high-throughput automated cryo-FIB milling^36^, increased the number of lamellae obtained per embryo (Figure S2B), and improved image contrast due to a ∼3-fold reduction in cryoprotectant concentration (Figure S2A). Using blastomeres from only 14 embryos, we resolved CPL filaments at 6.6 Å resolution using 180 tomograms (Figure 1E). Importantly, the CPL subtomogram density map obtained from individual blastomeres remained consistent with that resolved from intact embryos (Figure 1E) showing that the structure is not affected by blastomere separation. Moreover, the improved data quality reduced material requirements for high-resolution analysis, such that we estimated ∼80 embryos would be sufficient to achieve sub-6 Å reconstructions (Figure 1C).

To obtain a higher resolution structure of CPLs, we collected a large dataset of 1,153 tomograms from blastomeres derived from 66 embryos, yielding a consensus subtomogram average at 5.8 Å resolution (Figure S3). To account for conformational heterogeneity and flexibility between different regions of the filament, we performed particle symmetry expansion along with focused 3D classification and refinement, which further improved map quality to ∼4.7 Å resolution (Figure S3A and Table S1). We combined the resulting locally refined maps into a composite reconstruction (Figure 2A and Movie S1) with local resolution predominantly ranging from 4.5 to 6.0 Å (Figure S3B-C). Overall, the data demonstrates that our workflow enables robust in situ structure determination within mammalian embryo blastomeres while preserving the native architecture of macromolecular complexes.

**Figure 2.**
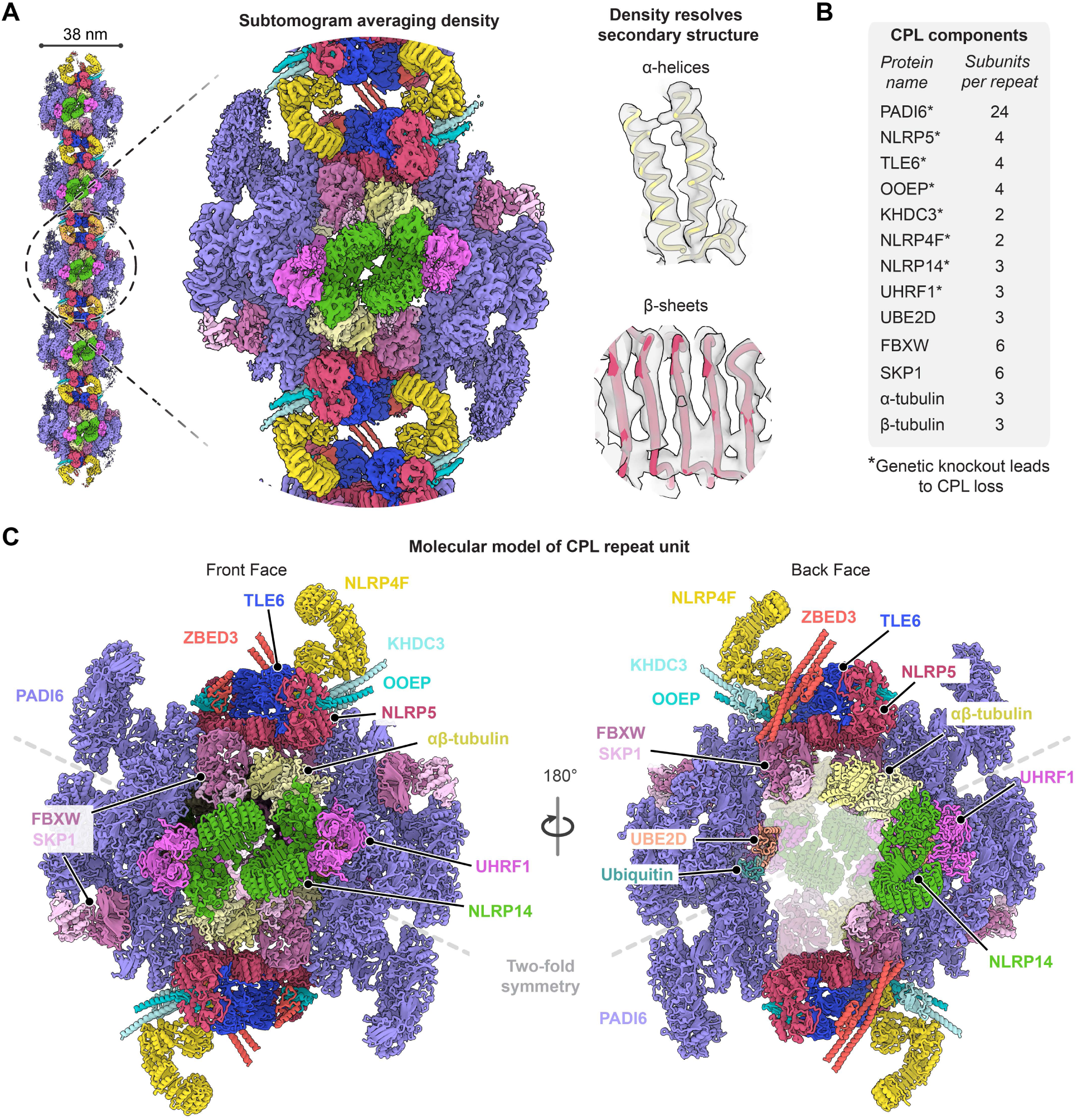
Structure of the CPL filaments. **(A)** Subtomogram averaging composite map of the CPL repeat unit. Examples of structural features resolved in the map are shown on the right. **(B)** Stable components identified within CPL filaments and their copy number per repeat. **(C)** Atomic model of the CPL repeat unit viewed from the front (left) and back (right), showing its overall organization. The overall protein arrangement exhibits a pseudo two-fold symmetry with the two symmetric halves separated by the grey dotted line.

### Molecular composition of CPL filaments in early mouse embryos

Our composite subtomogram averaging reconstruction displays well-resolved secondary structure features, including clear densities corresponding to α-helices and β-sheets (Figure 2A and S4A). This structural information was sufficient to assign all densities within the CPL filament repeat unit and build its molecular model. Each 38 nm repeat contains 14 distinct proteins present in multiple copies, resulting in a total of 71 subunits. These comprise PADI6, components of the SCMC, proteins involved in ubiquitination, NLRP14 as well as αβ-tubulin heterodimers (Figure 2B). Subunits on the front face are arranged with two-fold symmetry, whereas the symmetry on the back face is incomplete (Figure 2C). The combined molecular mass of each repeat unit is ∼4.5 MDa, placing CPLs among the largest known repeating units of a biological filament and, to our knowledge, second only to the axonemal 96 nm repeat^37^.

To identify the proteins within the CPL repeat, we adopted a candidate-based approach focussing on proteins implicated with CPLs. The rationale for each assignment is summarized in Table S2. We began with factors directly implicated in CPL formation, namely PADI6 and the SCMC, reasoning that these were most likely to be a part of the filaments. We identified 24 copies of PADI6 per repeat unit, making it the most abundant component of the CPL filament (Figure 2B and 2C). We further found a subset of SCMC components. Specifically, we detected the NLRP5-TLE6-OOEP complex along with KHDC3, ZBED3 and NLRP4F (Figure 2B and 2C). Thus, not all proteins classified as SCMC form part of the stable CPL repeat, indicating that a subset of SCMC proteins likely performs functions outside the assembled filament.

UHRF1 and NLRP14 are also important for CPL formation^11,12^. UHRF1 is an E3 ubiquitin ligase well-known for its role in maintenance of DNA methylation in the nucleus; however, in oocytes and early embryos it is predominantly cytoplasmic due to its association with CPLs^12,38^. NLRP14 has been implicated in stabilizing UHRF1 in the cytoplasm by protecting it from degradation^11^. Although both proteins had been proposed to be stored on CPLs, we found that UHRF1 and NLRP14 are integral components of the CPL repeat unit (Figure 2B and 2C), explaining why their loss leads to disappearance of CPLs.

The remaining un-assigned densities include additional components of the ubiquitination machinery. Density next to UHRF1 fit with its known E2 ubiquitin-conjugating enzyme UBE2D^39,40^ and ubiquitin (Figure S4B). In addition, there are multiple copies of a complex of a WD40 containing F-box protein and SKP1. F-box proteins mediate substrate recognition within SKP1-CUL1-F-box (SCF) Cullin-RING ligases^41–43^, whereas SKP1 functions as an adaptor that links F-box substrate receptors to the Cullin scaffold^42^. Our density shows a beta-hairpin insert into the WD40 domain that is consistent with the family of proteins homologous to the human FBXW12 protein (Figure S4A). In mice there are multiple possible proteins in this family including FBXW15, whose levels decrease upon knockout of PADI6^27^. However, at our resolution we are not able to discriminate between these homologs and so refer to these components as FBXW. We also identified an αβ-tubulin heterodimer as a structural element of the CPL repeat unit (Figure S4C and S4D), representing, to our knowledge, the only assembly outside of microtubules in which αβ-tubulin serves as a stable architectural component.

Notably, of these fourteen proteins, all except ZBED3, NLRP4F and FBXW are known to be enriched on CPLs by immunofluorescence^27^, supporting their localization to this structure. The assignments of the remaining three are further supported by previously reported interactions of ZBED3 and NLRP4F with the NLRP5-TLE6-OOEP complex^24,29^, as well as of FBXW with SKP1^42^. Importantly, ZBED and NLRP4F knockout mice are also known to lack CPLs in oocytes^23,24^.

Overall, our structure shows that CPL filaments are made of fourteen proteins and defines their architecture. It further provides the structural basis required to understand their role in early development.

### CPL repeats are built around PADI6 and NLRP5-TLE6-OOEP scaffold

Each CPL repeat unit is organized around a scaffold composed of PADI6 and the NLRP5-TLE6-OOEP complex (Figure 3A). All remaining CPL components assemble onto this scaffold.

**Figure 3.**
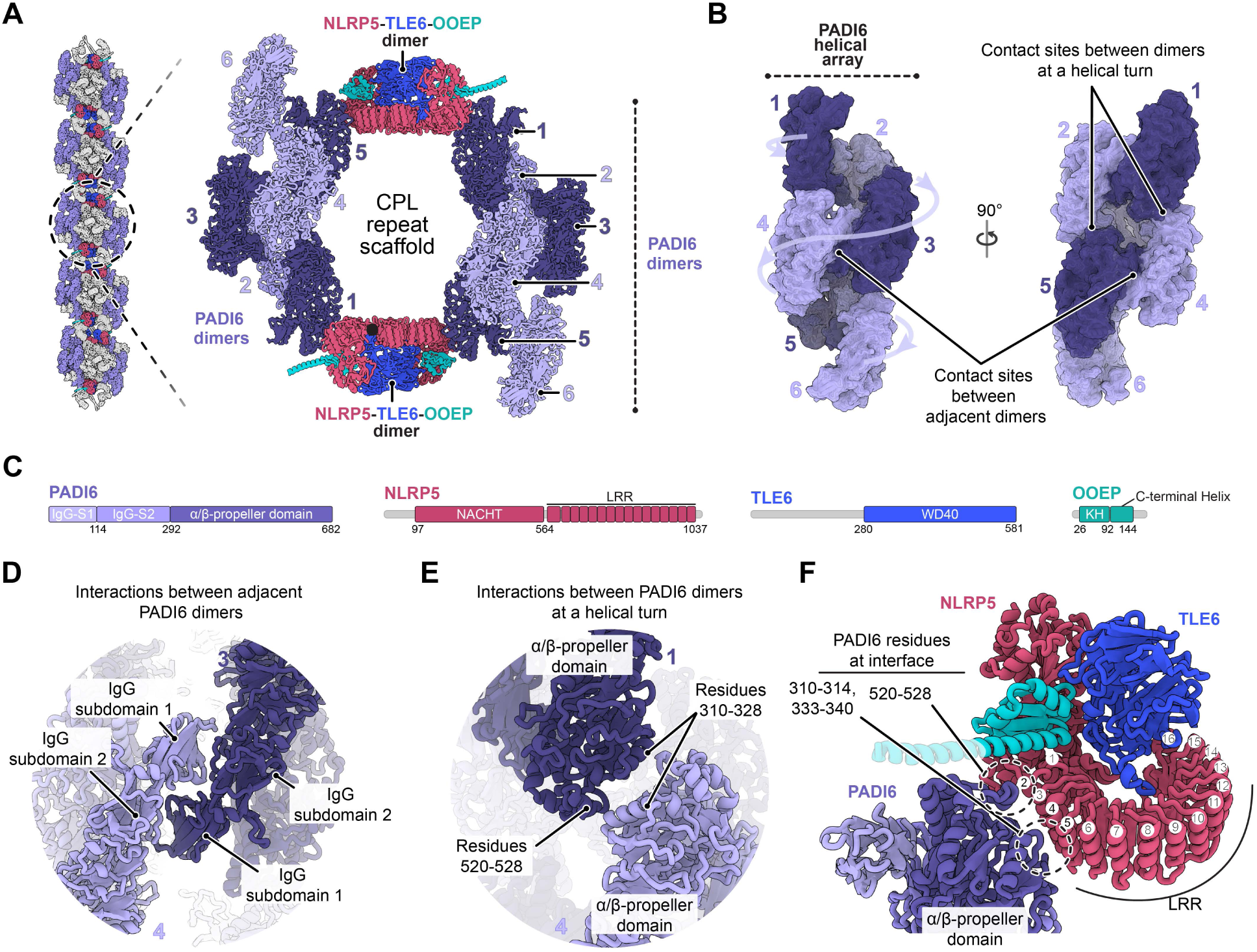
Organization of the proteins that scaffold CPL repeat units. **(A)** Arrangement of PADI6, NLRP5, TLE6 and OOEP within a single CPL repeat unit. The six PADI6 dimers are labelled 1-6 and shaded in two shades of lavender to distinguish them. Proteins in the core are hidden for clarity. **(B)** Helical array formed by PADI6 dimers, highlighting the two types of interdimer contacts. **(C)** Domain architecture of the CPL scaffold proteins. IgG-S, immunoglobulin subdomain; LRR, leucine-rich repeat; WD40, WD-repeat domain; KH, K-homology. **(D)** Close-up view of the IgG subdomains at the interface between adjacent PADI6 dimers. **(E)** PADI6 α/β-propeller domains at positions where the helix completes a turn (i.e., between dimers 1-4 and 2-5). **(F)** Interaction sites between NLRP5-TLE6-OOEP complex and PADI6.

PADI6 is a member of the peptidylarginine deiminase (PADI) enzyme family, which catalyzes protein citrullination, and, like other PADI enzymes, forms homodimers^44–46^. In our structure, PADI6 dimers assemble into a higher-order helical array (Figure 3B) that has not previously been described for proteins of the PADI family. Each array comprises six PADI6 dimers (Figure 3A-B), five of which are stably bound, whereas one remains flexibly attached (Figure S5A). Two such arrays form the lateral sides of the CPL scaffold (Figure 3A-B). PADI6 comprises two N-terminal immunoglobulin-like (Ig-like) subdomains and a C-terminal α/β-propeller domain (Figure 3C), both of which mediate interactions that stabilize the helical assembly (Figure 3D-E). The catalytic sites of PADI6, which face the cytoplasm in each CPL repeat, are occluded by neighbouring loop segments (Figure S5B-C). This conformation is consistent with structures of isolated PADI6 dimers (Figure S5C) and biochemical data which showed no detectable activity against canonical PADI substrates in vitro^44,45^. Therefore, our data suggests that PADI6 adopts a conformation incompatible with citrullination in situ and instead serves a structural role within CPLs.

The NLRP5-TLE6-OOEP complex is incorporated as a dimer within the CPL repeat. Two such dimers are present per repeat unit, with each bridging the two lateral PADI6 arrays (Figure 3A). Although previous in vitro studies detected both monomeric and dimeric forms^30^, we observe only the dimeric complex, indicating that this is the functionally relevant state within CPLs. NLRP5 comprises an N-terminal NACHT domain and a C-terminal leucine-rich repeat (LRR) domain (Figure 3C). The NACHT region engages TLE6 and OOEP, whereas the LRR domain mediates direct interactions with PADI6 (Figures 3F). This arrangement establishes a direct structural linkage between PADI6 and the NLRP5-TLE6-OOEP complex. In contrast to previous proposals that these factors act as independent modules^27,47^, our findings provide a structural explanation for the reported interactions between them and show that they assemble into a unified scaffold.

### SCMC proteins link CPL repeat units to form filaments

Adjacent CPL repeat units are linked by three bridging factors, KHDC3, ZBED3 and NLRP4F, which connect neighbouring repeats through interactions with the NLRP5-TLE6-OOEP complex (Figure 4A and S6). KHDC3 binds the front-facing side of each NLRP5-TLE6-OOEP dimer (Figure 4B-C), whereas ZBED3 engages the opposite side at the back face of the filament (Figure 4D). ZBED3 contains an N-terminal zinc-finger (ZnF) domain and a C-terminal helical region (Figure 4B). The ZnF domains of two ZBED3 molecules bind TLE6. The C-terminal helical regions of the two ZBED3 molecules from adjacent repeat units further associate to form a tetrameric helical bundle that directly links neighbouring repeats (Figure 4D). Together, KHDC3 and ZBED3 create two distinct binding surfaces on the NLRP5-TLE6-OOEP dimer (Figure 4E). The KHDC3-associated side accommodates the N-terminal domains of NLRP4F (Figure 4F), whereas the ZBED3-associated side binds the C-terminal LRR domain of NLRP4F (Figure 4G). In this arrangement, two NLRP4F molecules span neighbouring repeat units in an antiparallel orientation (Figure 4A).

**Figure 4.**
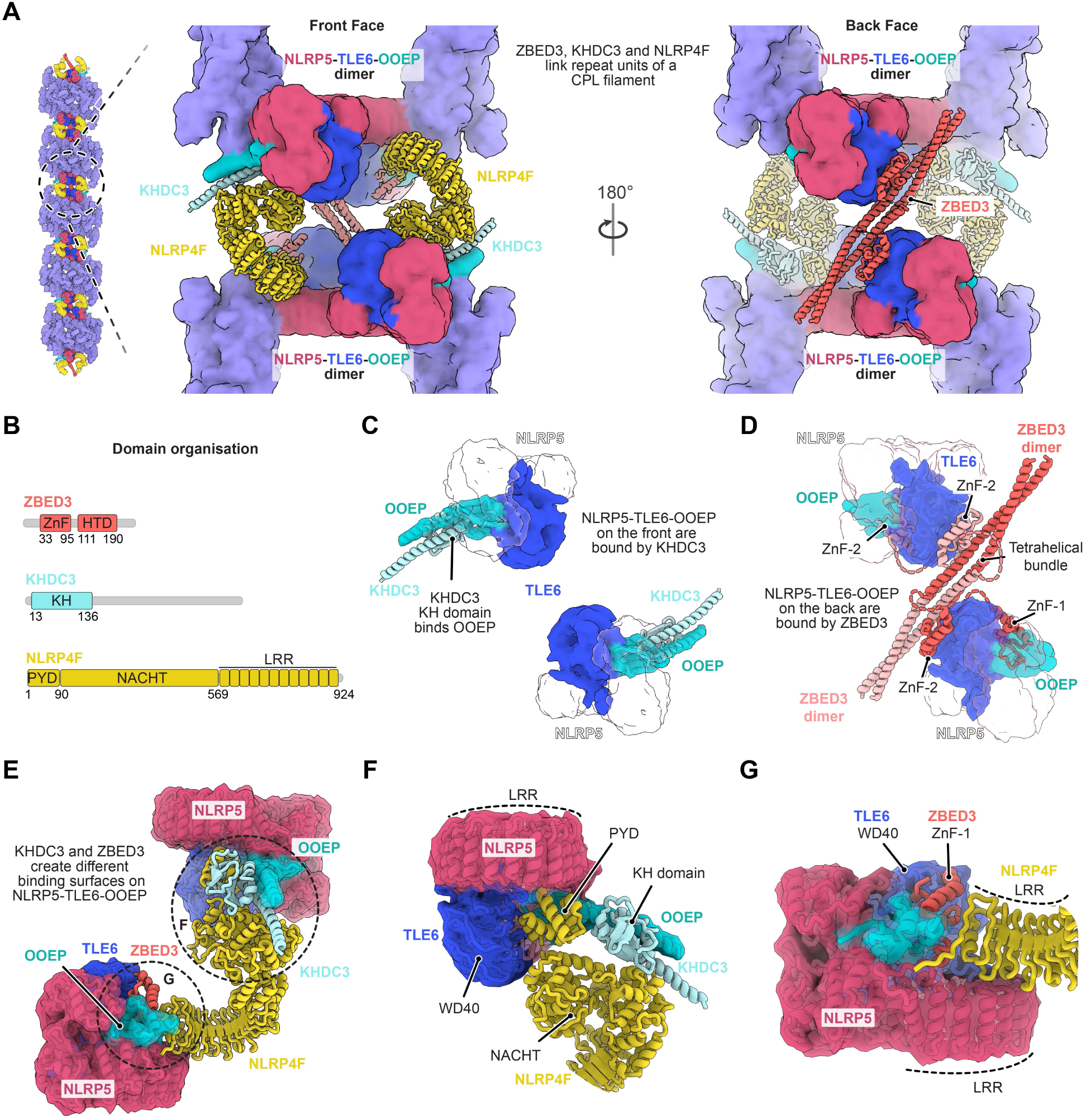
Molecular basis for the formation of CPL filaments. **(A)** Front (left) and back (right) faces of the segment between two CPL repeat units. KHDC3, ZBED3 and NLRP4F cooperate to link adjacent repeats. **(B)** Domain architecture of KHDC3, ZBED3 and NLRP4F. KH, K-homology; ZnF, BED type Zinc finger; HTD, Helical Tetramerization Domain; PYD, Pyrin domain; and leucine-rich repeat (LRR). **(C, D)** The interactions of KHDC3 (C) and ZBED3 (D) with the NLRP5-TLE6-OOEP complexes. The ZnF domain of ZBED3 that contacts both the KH domain of OOEP and the WD40 domain of TLE6 (labelled as ZnF-1) is stably bound, whereas the ZnF domain that interacts only with the TLE6 WD40 domain (labelled as ZnF-2 in panel D) is more flexibly associated (Figure S6). ZBED3 from adjacent repeats forms a four-helix bundle that links adjacent repeats. **(E)** Contacts made by NLRP4F while linking two adjacent repeat units. **(F)** Interaction of the N-terminal PYD and NACHT domain of NLRP4F with KHDC3 **(G)** Interaction of C-terminal LRR of NLRP4F with ZBED3 bound NLRP5-TLE6-OOEP complex.

Genetic data shows that deletion of any of the three bridging factors results in the loss of CPLs ^23,24,26^. The phenotype observed in *Nlrp4f* mutants suggests NLRP4F plays a major role in holding the CPL repeats together and that the connection mediated by ZBED3 alone can’t compensate for its function. The effect of *Khdc3* and *Zbed3* mutants is explained by their disruption of the binding sites for NLRP4F on the NLRP5-TLE6-OOEP dimers. Thus, all three bridging factors cooperate to link neighbouring repeat units and promote CPL filament formation.

Overall, we show how different components of the SCMC interact within CPLs and define their stoichiometry. The SCMC was originally described based on enrichment of its components beneath the oocyte cortex^7^, in contrast to CPLs, which are distributed throughout the cytoplasm^3–5^. However, contrary observations^9,27^ showing a uniform cytoplasmic distribution of SCMC components and their colocalization with CPLs have raised questions about the relationship between these two assemblies. Our data now provides direct evidence that the key components of the SCMC are part of the CPL filament.

### CPLs stably incorporate ubiquitination proteins and tubulin

Our structure reveals that CPLs contain multiple FBXW-SKP1 complexes (Figure 5A) together with a distinct assembly composed of UHRF1, UBE2D, NLRP14, and an αβ-tubulin heterodimer (Figure 5B). These are positioned within the central region and complete the architecture of the repeat unit (Figure 5C).

**Figure 5:**
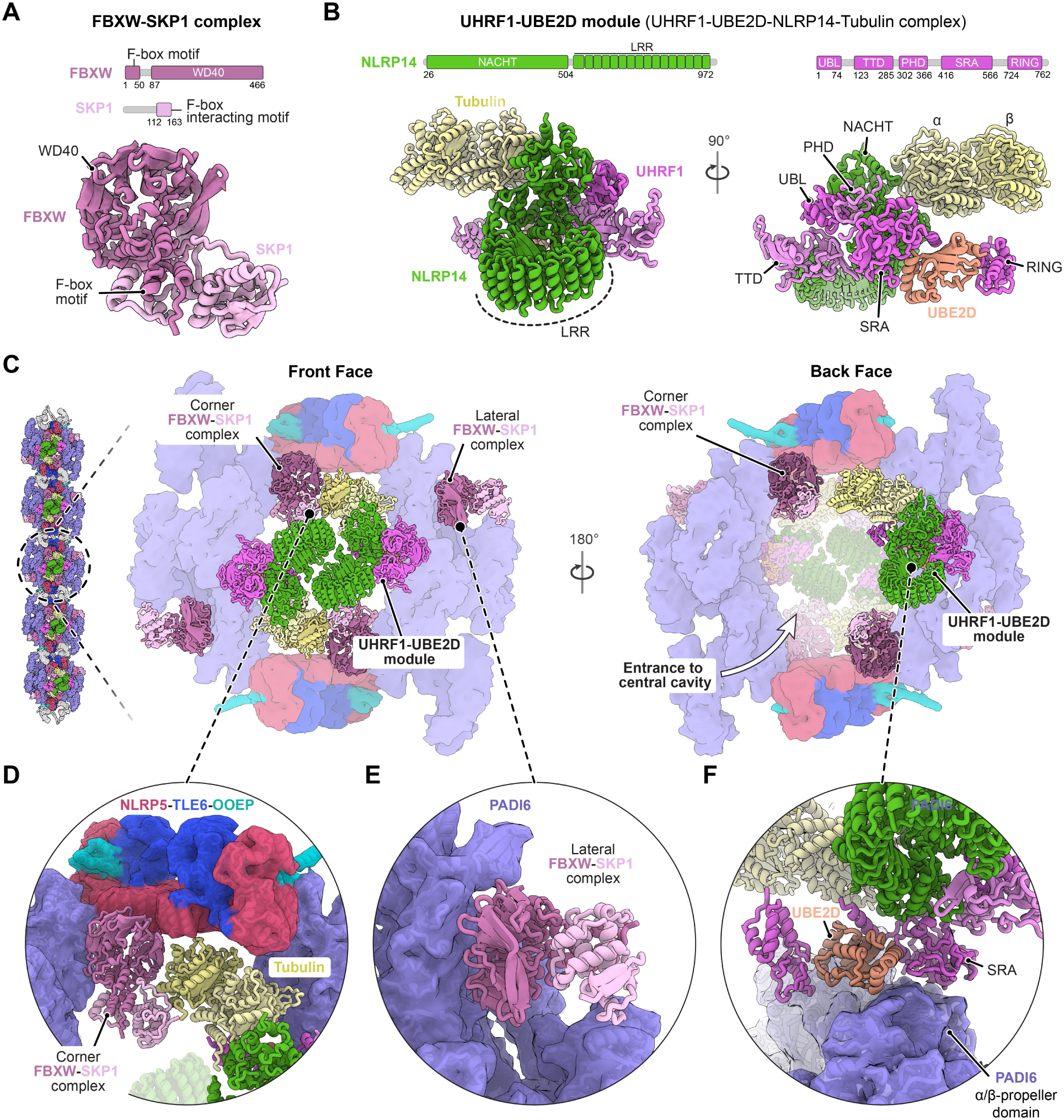
Organization of the proteins from the ubiquitination pathway within CPL repeat units. **(A)** Domain organization and structure of the FBXW-SKP1 complexes. **(B)** Domain organization and architecture of the E2-E3 assembly composed of UHRF1-UBE2D-NLRP14-αβ-tubulin. **(C)** Arrangement of FBXW-SKP1 and E2-E3 assemblies within a single CPL repeat unit, shown are front and back faces of the filament to highlight differences in protein composition and placement. **(D-F)** Zoom-in showing the interaction of corner FBXW-SKP1 complex with tubulin of front UHRF1-UBE2D modules (D), binding of the lateral FBXW-SKP1 to the PADI6 helical array (E) and interaction of UHRF1 SRA domain and UBE2D with PADI6 (F).

The FBXW-SKP1 complex is formed through the interaction of the F-box motif of the FBXW with SKP1 (Figure 5A), consistent with other F-box-SKP1 assemblies^42^. Six of these complexes are anchored directly to the PADI6 and NLRP5-TLE6-OOEP scaffold (Figure 5C). Four occupy corner positions between NLRP5 and PADI6 (Figure 5D), while two bind along the outer surface of the PADI6 arrays (Figure 5E). These interactions are mediated by the FBXW through their WD40 domain. Notably, the canonical WD40 substrate-binding side used by other F-box proteins^42^ is involved in binding the scaffold (Figure S7A). The FBXW-SKP1 complexes engage in three distinct types of interfaces: the front-facing corner complexes contact β-tubulin (Figure 5D and S7B), the back-facing corner complexes contact ZBED3 (Figure S6), and the lateral complexes mediate the inter-filament interactions described later. Together, these observations indicate that FBXW-SKP1 complexes play multiple structural roles within CPLs.

A second assembly, hereafter referred to as the UHRF1-UBE2D module, is organized around NLRP14 (Figure 5B). In this module, NLRP14 simultaneously binds α-tubulin and engages multiple N-terminal domains of UHRF1, namely, the ubiquitin-like (UBL), tandem Tudor (TTD), plant homeodomain (PHD), and SET- and RING-associated (SRA) domains. On the other hand, UBE2D associates with the SRA, its proximal helix and the C-terminal RING domain of UHRF1 (Figure 5B). This contrasts with a recent study where no RING domain was observed bound to UBE2D and only the SRA domain of UHRF1 was involved in binding UBE2D^39^. Our structure further shows that, in the context of the CPL filament, UHRF1 and UBE2D are incorporated into a larger assembly together with NLRP14 and tubulin. Three such modules are present per repeat: two are positioned symmetrically on the front face and are stabilized by contacts with PADI6 and adjacent FBXW-SKP1 complexes (Figure 5C-D), whereas the third occupies the back face and is anchored primarily through interactions with PADI6 (Figure 5C and 5F).

The FBXW-SKP1 complexes and UHRF1-UBE2D modules not only form integral components of the repeat unit but also mediate higher-order organization of filaments into lattices. In all cases, inter-filament connections are established by FBXW-SKP1 and PADI6 on the outer surface of one filament engaging NLRP14 and N-terminal domains of UHRF1 on the back face UHRF1-UBE2D module of a neighbouring filament (Figures 6A and Figure S8A-B). Multiple copies of these paired interactions cooperate to generate stable inter-filament linkages (Figure 6B). To define how these contacts give rise to lattice organization, we performed 3D classification using a larger particle box that included neighbouring filaments (Figure S8C). This analysis showed that lattice assembly is heterogeneous, spanning a range of attachment patterns from filaments with no stably attached neighbours to filaments engaged with up to three neighbours (Figure 6C and S8C). These contacts occur at defined attachment points, with each filament forming up to two lateral interactions and one back-face interaction (Figure 6D).

**Figure 6.**
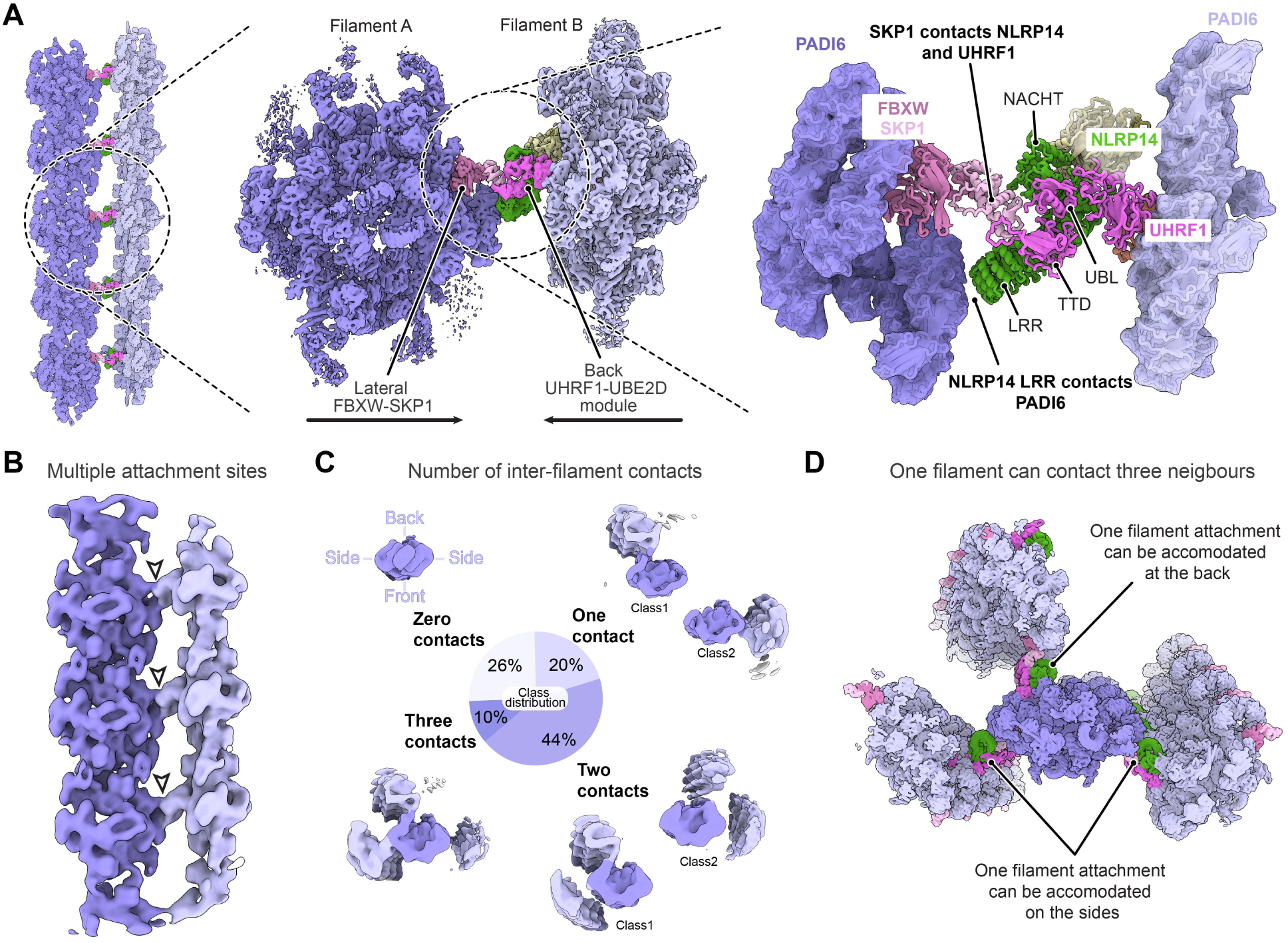
Molecular basis for organisation of CPL filaments into lattices. **(A)** Inter-filament linkages mediated by the FBXW-SKP1 and PADI6 on the side of one filament contacting the UHRF1-UBE2D modules at the back face of an adjacent filament. **(B)** Multiple adjacent inter-filament attachment points are shown along the inter-filament interface. **(C)** Distribution of 3D classes showing the number of stable inter-filament contacts observed within CPLs. Top views of the representative classes are displayed. **(D)** The three sites that can be employed for interfilament contacts are depicted.

### The UHRF1-UBE2D module can be charged with ubiquitin

The importance of the CPL constituents identified in the structure and their higher order assembly is indicated by genetic loss-of-function studies^7–12,23,24,26^. We find proteins whose absence results in embryonic lethality or infertility comprise the CPL scaffold (PADI6 and NLRP5-TLE6-OOEP) or reside within it (UHRF1, UBE2D, NLRP14 and SKP1). In contrast, deletion of bridging factors required for filament assembly (KHDC3, ZBED3, and NLRP4F) permits viable progeny despite CPL loss. Together, these findings demonstrate that filament formation is not essential for development, whereas the scaffold and the assemblies within it form the functional unit of CPLs required for developmental success. We therefore refer to this functional unit as the CPL core complex (Figure S9).

A distinctive feature of the CPL core complex is the arrangement of UHRF1-UBE2D modules, which generates a large central cavity (∼2000 nm³) within each repeat (Figure 5C). This cavity contains all three UBE2D molecules (Figure 7A) and remains accessible to cytoplasmic proteins through an ∼15 nm opening on the back face of the filament.

**Figure 7:**
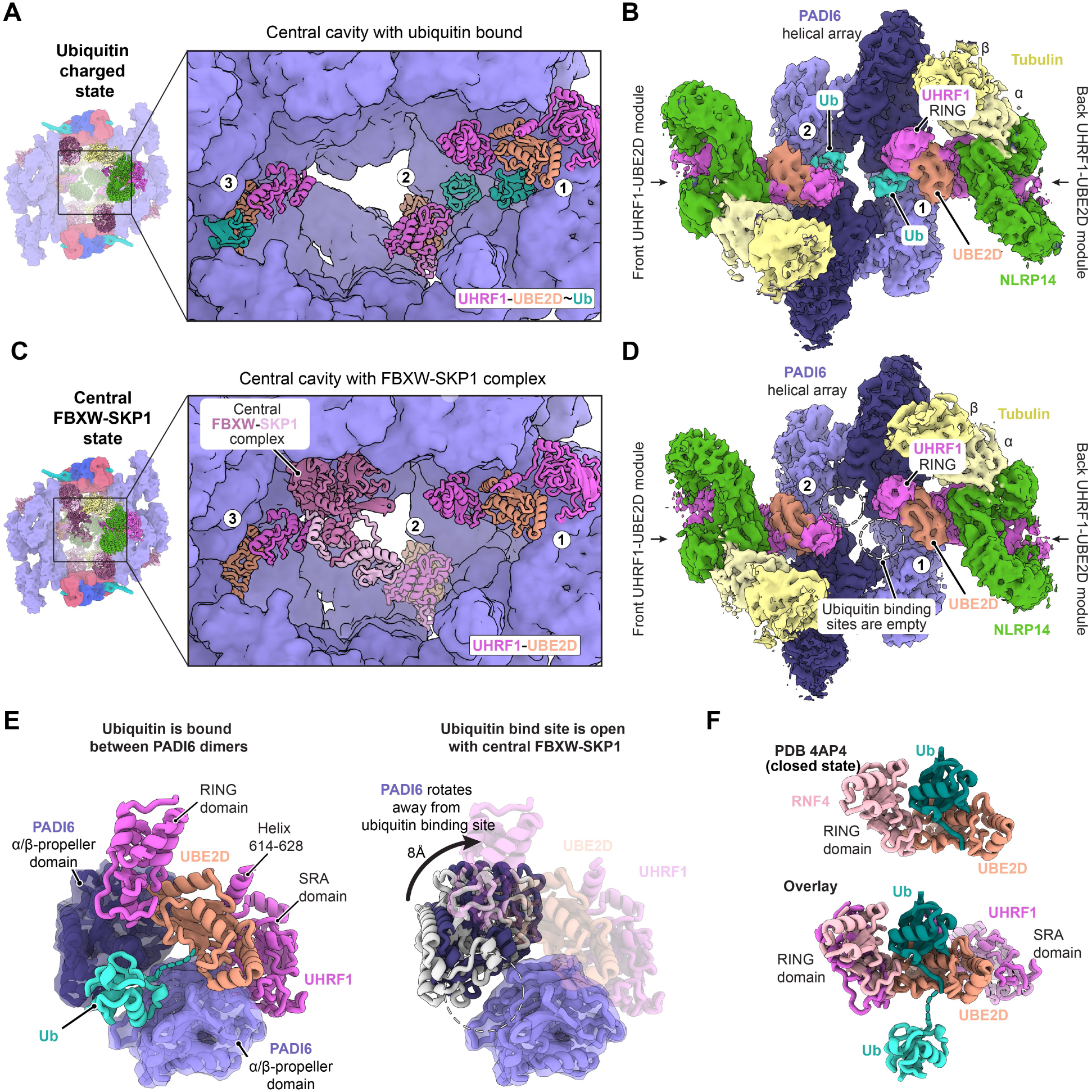
Central cavity within CPL repeats contains ubiquitin-charged UBE2D. **(A)** Central cavity within a CPL repeat unit viewed from the back face, showing the arrangement of the three UHRF1-UBE2D (RING-E2) modules and bound ubiquitin moieties. **(B)** Subtomogram averaging density corresponding to the UHRF1-UBE2D modules labelled 1 and 2 in A. **(C)** Central cavity within a CPL repeat unit in the state containing a central FBXW-SKP1 complex. **(D)** Subtomogram averaging density corresponding to the UHRF1-UBE2D modules labelled 1 and 2 in C in the presence of the additional FBXW-SKP1 complex within the cavity. **(E)** Comparison of the ubiquitin-bound configuration and the central FBXW-SKP1 state. In the ubiquitin-charged state, ubiquitin is stabilized between the α/β-propeller domains of two PADI6 dimers. In the FBXW-SKP1 state, rearrangements of the PADI6 helical array open the ubiquitin-binding pocket. **(F)** Structure of UBE2D bound to a RING domain with ubiquitin in the closed conformation (PDB-4AP4)^65^. In this configuration, ubiquitin is positioned between the RING domain and the E2 enzyme, representing the catalytically competent state. Overlay with the UHRF1-UBE2D-ubiquitin configuration observed in CPLs shows that ubiquitin adopts an open conformation relative to the closed active state.

Using 3D classification, we resolved two distinct structural states of this cavity (Figure 7A-D). In the first state, density corresponding to ubiquitin (Ub) is observed adjacent to all three UBE2D molecules (Figure 7B and S10A). The distance between the ubiquitin and the UBE2D catalytic cysteine is consistent with a covalently linked E2∼Ub conjugate^48–50^. The ubiquitin moiety is stabilized by PADI6 and lodged between the α/β-propeller domains of two PADI6 dimers (Figure 7E), revealing an unexpected role for PADI6 in stabilizing ubiquitin bound to an E2 enzyme. In this configuration, the E2∼Ub is held in an open conformation that is associated with low activity for ubiquitin transfer^51^. This contrasts with the catalytically active state which is promoted by RING domains through stabilization of the closed E2∼Ub conformation, in which ubiquitin engages both the E2 and the RING domain^48–50^ (Figure 7F).

In the second state of the cavity, there is an additional centrally positioned FBXW-SKP1 complex which is bound to a corner FBXW-SKP1 complex and the β-tubulin of the back face UHRF1-UBE2D module (Figure 7C, S7B and S10B). Strikingly, however, in this state there is no ubiquitin density present adjacent to the UBE2D subunits (Figure 7D). Structural comparison with the ubiquitin-bound state shows that presence of the central FBXW-SKP1 complex is accompanied with rearrangements within the CPL core complex. The back UHRF1-UBE2D module moves inwards so that its tubulin contacts the central FBXW-SKP1. This results in the rearrangement of the PADI6 helical arrays, and the opening of the ubiquitin binding pockets (Figure 7E and Movie S2).

The opening of these ubiquitin-binding pockets likely explains the absence of ubiquitin density when the central FBXW-SKP1 complex is bound. Ubiquitin may remain tethered to UBE2D but be too flexible to visualize. Such a state would permit ubiquitin to engage the RING domain and UBE2D in the catalytically active closed conformation, thereby making it available for transfer to substrates. Alternatively, given the short half-life of E2∼Ub conjugates bound to an E3 ligase^51^, ubiquitin may already have been transferred onto a substrate. Together, these two structural states reveal that the CPL core complex can accommodate ubiquitin-charged UBE2D molecules and undergo conformational rearrangements that are consistent with regulated ubiquitin transfer.

## Discussion

This study provides a general strategy for structural analysis of mammalian embryos at resolutions sufficient for protein identification and visualizing different states of protein complexes present in situ. By separating 6/8-cell stage mouse embryos into individual blastomeres prior to vitrification, we generated samples compatible with automated cryo-FIB milling. The resulting improvement in sample quality enabled reconstructions at 6.6 Å resolution using only 14 embryos. Such low material requirements make this approach particularly attractive for studies of human embryos donated for research and for investigating intracellular organization in samples derived from patients with infertility. Although demonstrated here at the 6/8-cell stage, the workflow is readily applicable to other stages of preimplantation development, providing a broadly applicable approach for structural investigation of macromolecular assemblies in early mammalian embryos.

Despite occupying 10% of the cytoplasmic volume of early embryos^27^, the function of CPLs is currently unclear. One proposed role is protein storage^27^. However, all density in our subtomogram averages can be accounted for by the assigned proteins, raising the question of whether and how the hundreds of maternal proteins proposed to be stored on CPLs^27,28^ associate with the filamentous structure. For some candidate stored factors, such as 14-3-3 proteins and the SCMC interactor NLRP2, binding within the lattice appears unlikely, as their reported binding sites on the NLRP5-TLE6-OOEP complex^29^ would clash in the context of the CPL structure (Figure S11A-B). Other factors may associate flexibly or sparsely with the CPLs meaning we would not resolve them in our averaged electron density. One site for such interaction is the C-terminal region of KHDC3 which is known to bind SPIN1^52^ and is disordered in our structure. It is also possible that the CPLs are specifically sequestering tubulin and components of the ubiquitination machinery, namely the FBXW-SKP1 and the UHRF1-UBE2D-NLRP14-tubulin complexes. This would link CPLs with a wide range of processes, given that tubulin is important for microtubule cytoskeleton^53^, FBXW-SKP1 complexes are linked to the SCF-type ubiquitin ligase machinery^41,42^, UBE2D is a promiscuous E2 enzyme^54,55^, and UHRF1 has a key role in the maintenance of DNA methylation^56,57^. Such a mechanism could help explain the diverse defects observed upon CPL loss^10,22–28^. However, because these proteins are stable components of the lattice even until the 8-cell stage embryos, how their release would be regulated, or at what stage of development this might occur is unclear.

In contrast to a purely storage role, our in situ structure suggests that CPLs are directly involved in ubiquitination. We find that all three UBE2D molecules within a CPL repeat unit can adopt a ubiquitin-charged state. These UBE2D∼ubiquitin conjugates are held in a catalytically unfavourable open conformation through interactions between ubiquitin and PADI6, likely limiting ubiquitin transfer onto non-specific targets. Strikingly, in the presence of a central FBXW-SKP1 complex, the ubiquitin-binding sites on PADI6 rearrange, and the ubiquitin density is no longer resolved, suggesting that it becomes available for transfer. In our structure, the UBE2Ds are bound to UHRF1 RING domains. Upon the release of ubiquitin from PADI6 these domains are positioned to promote a catalytically active closed state of the UBE2D∼ubiquitin conjugate. Together, the architecture of CPLs indicate they possess the important features required for ubiquitin transfer: ubiquitin-charged E2 enzymes, a mechanism for regulating ubiquitin availability, and RING domains positioned to promote the catalytically active state.

Interestingly, UBE2D binding by UHRF1 involves a helix that contacts the backside surface of the E2 enzyme. Comparable backside interactions have been shown to promote monoubiquitination in other RING E3 ligases^58,59^ (Figure S11C). This is also consistent with the established nuclear functions of UHRF1, where it mediates monoubiquitination of substrates such as PAF15 and histone H3^56,60^.

Within CPL filaments, the canonical substrate-binding domains of UHRF1 (TTD, PHD and SRA domains)^61,62^ all contact NLRP14, suggesting that they are not available for conventional substrate recognition. However, the architecture of the central cavity places the UHRF1-UBE2D modules adjacent to multiple FBXW-SKP1 complexes, whose homologs are best known as substrate receptors in SCF Cullin-RING ligases^41–43^. We speculate that these FBXW-SKP1 complexes are repurposed here for a similar role in substrate recruitment. In particular, the central FBXW-SKP1 complex could serve a dual function by both recruiting a bona fide substrate and triggering release of ubiquitin from the PADI6-bound state, thereby promoting ubiquitination only when substrate is properly engaged. Because ubiquitin-dependent regulation can influence protein activity, localization and stability^63,64^, CPLs acting as sites of ubiquitination could also help explain the diverse defects observed upon CPL loss.

Together, based on our observations we propose that CPLs act as a giant E2-E3 ubiquitin ligase assemblies that organize multiple UHRF1-UBE2D modules within a central reaction chamber. Upon binding of the central FBXW-SKP1 complex, structural rearrangements promote the transition from a restrained ubiquitin-bound state to a catalytically permissive state allowing ubiquitin transfer onto a substrate.

## Methods

### Mouse embryo collection

Zygotes were collected from the oviducts of ∼8-12-week-old timed-mated CD-1 mice (Charles River), and granulosa cells were removed by brief (∼5 s) treatment with hyaluronidase (Sigma-Aldrich; #H4272, at 1 mg/ml, stored at −20°C). Embryos were then washed and cultured at 37°C under 5% CO₂. All procedures were performed in EmbryoMAX® Advanced KSOM medium (Millipore; #MR-101-D). All mice were maintained at 21-23°C, 45-65% humidity, under a 12 h light/dark cycle (6:30 −18:30).

### Mouse embryo blastomere separation and reassociation

Zona pellucida of 6/8-cell mouse embryos was removed by brief (∼10 s) exposure to acidic Tyrode’s solution (Sigma-Aldrich; T1788), followed by several washes in EmbryoMAX® Advanced KSOM medium (Millipore; #MR-101-D). Embryos were subsequently incubated in 0.5% trypsin, 1 mM EDTA solution for up to 2 min at 37°C (prepared fresh in PBS from 2.5% trypsin, Gibco 15090046, and 0.5 M EDTA pH 8.0, Invitrogen 15575-038), then transferred to 4% BSA, 0.5 mM EDTA solution to inactivate trypsin (prepared fresh in PBS from 30% BSA, Sigma A7284, and 0.5 M EDTA pH 8.0, Invitrogen 15575-038). Mechanical dissociation of blastomeres was performed in the 4% BSA, 0.5 mM EDTA solution using a transfer pipette with a tip of 50 µm diameter attached to an EZ-Grip denudation pipettor. Disaggregated blastomeres were washed through and finally cultured in 5 µl droplets of pre-equilibrated EmbryoMAX® Advanced KSOM medium at 37°C under 5% CO₂. Small indentations were created in 5 µl culture droplets using a custom-fabricated stainless-steel needle to facilitate blastomere reaggregation.

### Immunofluorescence

Mouse embryos were fixed in 4% methanol-free formaldehyde (Thermo Scientific; #28906) in phosphate-buffered saline (PBS) for 20 min at room temperature. Fixed embryos were permeabilized in PBS containing 0.2% Triton X-100 for 10 min at room temperature, then blocked in 1% BSA (Fisher BioReagents; BP1605-100) in PBS overnight at 4°C. Primary antibodies were incubated overnight at 4°C. The following primary antibodies were used: anti-SOX2 (Invitrogen; #14-9811-82, 1:200) and anti-GATA3 (Abcam; ab199428, 1:200). Secondary antibodies and DAPI (Vector Laboratories; #H-1200) were applied for 1 h at room temperature. Secondary antibodies were Donkey anti-Rabbit IgG (H+L), Alexa Fluor 488 (Invitrogen; #A-21206) and Donkey anti-Rat IgG (H+L), Alexa Fluor Plus 555 (Invitrogen; #A48270), both used at 1:400 dilution in 1% BSA in PBS.

### Confocal imaging

Confocal imaging was performed in PBS with 3 mg/ml polyvinylpyrrolidone (Sigma-Aldrich; #P0930) under paraffin oil in a 35 mm dish with a #1.5 glass coverslip. Images were acquired with an SP8 confocal laser scanning microscope (Leica) using a plan apochromatic 40x 1.10 NA wa-ter-immersion objective.

### Grid preparation

6/8-cell mouse embryos were sequentially incubated in EmbryoMAX® Advanced KSOM medium supplemented with increasing concentrations of DMSO and ethylene glycol to aid vitrification^27^: 1.75% each for 3 min, 2.33% each for 3 min, 3.5% each for 6 min, and 7.5% each for 3 min. Separated blastomeres from 6/8-cell mouse embryos were incubated in a similar manner but only until the second step which contained 2.33% of each solution. Intact embryos or separated blastomeres were then applied onto freshly glow-discharged Quantifoil R2/2 200-square-mesh gold grids (Quantifoil) along with ∼2-3 μl of the final media solution. The grid was then placed in an EM GP2 plunge freezer (Leica) at 95 % humidity and 25°C, and blotted for 6 s before being plunged into liquid ethane.

### Cryo-FIB lamella preparation

Grids were clipped into Autogrids (Thermo Fisher Scientific) and subjected to automated lamella preparation using an Aquilos 2 FIB-SEM (Thermo Fisher Scientific). The grids were first sputter-coated with an inorganic platinum layer for 30 s followed by a coating of organometallic platinum using a gas injection system (GIS) for 40 s and an additional sputter-coat of an inorganic platinum layer for 30 s. Next, grids were surveyed using Maps software (Thermo Fisher Scientific) for lamella site identification. For separated blastomeres, this was followed by automated lamella preparation using AutoTEM Cryo (Thermo Fisher Scientific). Lamella were milled at an angle of 12° with a 30 kV ion beam in 4 steps: 1) 1 nA, gap 3 μm with stress relief cuts, 2) 0.5 nA, gap 2 μm, 3) 0.3 nA, gap 1.4 μm, 4) 0.1 nA, gap 0.7 μm. Lamellae were finally polished at 30-50 pA with a gap set to 100 nm. For intact embryos, lamellae were prepared manually. Material in front and back of the area of interest of ∼30-40 μm were milled away at an angle of 90°. Lamella were then milled using a similar stepwise milling protocol as above along with stress-relief notches or cuts to minimize lamella deformation.

### Cryo-ET data acquisition

For intact embryo samples, tilt-series data were collected using a Titan Krios G3i transmission electron microscope (Thermo Fisher Scientific) operating at 300 kV equipped with a K3 detector and an energy filter with a 20 eV slit size (Gatan) and 100 μm objective aperture. Dose-symmetric tilt series in 3° steps were acquired using Tomography 5 software (Thermo Fisher Scientific) with a tilt span of ± 48° around the milling angle. Target focus for each tilt-series was set in the range of −2 µm to −4.5 µm. Movies for each tilt were acquired at a magnification of 53,000x in tiff format with 6 frames at a pixel size of 1.63 Å and a nominal dose of 3.5 e-/Å^2^.

For separated blastomere samples, tilt-series data were collected using a Titan Krios G4 transmission electron microscope operating at 300 kV equipped with a Selectris X energy filter with a slit set to 10 eV, 100 μm objective aperture and a Falcon 4i direct electron detector (Thermo Fisher Scientific). Dose-symmetric tilt series in 3° steps were acquired using Tomography 5 software (Thermo Fisher Scientific) with a tilt span of ± 48° around the milling angle. Target focus for each tilt-series was set in the range of −2 µm to −4.5 µm. Movies for each tilt were acquired at a magnification of 64,000x (180 movies at 1.91 Å/pix) for the 14 embryo dataset or 105,000x (1153 movies at 1.185 Å/pix) for the larger 66 embryo dataset in tiff format with 6-8 frames at a nominal dose of 3.5 e-/Å^2^.

### Cryo-ET data processing and subtomogram averaging

All data processing was performed in RELION-5^66^ unless specified otherwise and is described in Figure S3. The processing for both the intact embryo samples and separated blastomeres were performed in a similar manner and here we explain the processing steps in the context of the latter sample dataset collected at the pixel size of 1.185 Å/pix. Global motion correction and dose-weighting were performed in RELION using MotionCor2^67^ with a B-factor of 150 and 5x5 patches. The power spectrum of aligned frames was used for contrast transfer function estimation by CTFFIND4.1^68^. Tilt series alignment was then performed using Aretomo2^69^ followed by tomogram generation in RELION at a pixel size of 10 Å. CPL filaments were first picked using PyTOM^70^ using EMDB-16458^27^ filtered to 40 Å as a reference to get an initial model and then this initial model was used for particle picking using PyTOM. Particles were extracted with a box size of 128 pixels at a pixel size of 4.74 Å followed by 3D refinement with the initial model filtered to 60 Å as a reference. 3D classification without alignment was performed and 134,387 particles that belonged to classes displaying clear features were selected followed by another round of 3D refinement. This was followed by CTF refinement and Bayesian polishing at pixel size of 2.37 Å and box size of 256 pixels resulting in a structure at 8.7 Å resolution. This was followed by two rounds of particle extraction with a box size of 256 pixels at a pixel size of 2.37 Å followed by 3D refinement (with C2 symmetry), CTF refinement and Bayesian polishing. The particles were then re-extracted at a pixel size of 1.185 Å and box size of 420 pixels followed by 3D refinement with C2 symmetry to obtain a 5.8 Å structure. Given the pseudo C2 symmetry of the CPL repeat unit, particles were symmetry-expanded in RELION. A subsequent 3D refinement in C1 symmetry, using a mask encompassing the asymmetric unit, improved the resolution to 4.7 Å. Focused 3D classifications (without alignment) and refinements were then performed on distinct structural regions. To resolve the inter-repeat bridging proteins, symmetry-expanded particles were recentered on ZBED3 tetrahelical bundle and extracted with a 180-pixel box at 2.37 Å/pixel. These were subjected to focused 3D classification and refinement using a mask that included the bridging proteins and the NLRP5-TLE6-OOEP dimers on opposing CPL repeats. To examine the inter-filament interaction between lateral FBXW-SKP1 and NLRP14-UHRF1, symmetry-expanded particles were recentered on lateral FBXW-SKP1 and extracted with a 180-pixel box at 2.37 Å/pixel, followed by focused 3D classification and refinement. For resolving the ubiquitin bound and central FBXW-SKP1 bound states, a mask around the back UHRF1-UBE2D module was used for 3D classification and refinement was performed. This was followed by removal of duplicate particles for each class and a 3D refinement with C1 symmetry using a mask for the full CPL repeat unit.

For visualizing the landscape of inter-filament contacts, the particles from the dataset collected on blastomeres from 14 embryos was used at it was at a larger pixel size which helped with 3D classification as it included more surrounding filaments than would be possible with the lower pixel size dataset. All particles contributing to the consensus structure were used and were re-extracted with a box size of 200 pixels at a pixel size of 7.64 Å followed by 3D classification without alignment and no mask.

A composite map for a CPL repeat unit was generated by merging maps from all the local refinements in UCSF ChimeraX^71^ using the “Volume max” function.

### Model building and refinement

The starting models used for all proteins, details of the residues built and the rationale for density assignment is summarized in Table S2. The protein complexes, proteins or protein domains were placed into the STA maps using UCSF ChimeraX guided by secondary structure features. The model then went through multiple cycles of restrained flexible fitting using ISOLDE^72^ (distance restraints ranging from 3-8 Å were imposed on starting models) followed by user-guided refinement in COOT^73^. Final model refinement and model validation was performed in PHENIX^74^. All refinement statistics can be found in Table S1.

### Data Visualization

Tomograms were denoised using cryoCARE^75^. Membranes were segmented using MemBrain v2^76^. Ribosomes and microtubules were picked using PyTOM for tomogram segmentation. ArtiaX plugin^77^ within UCSF Chimerax was used for mapping back structures to their coordinates in the original tomogram as displayed in Figure 1B. All structural figures were rendered in UCSF Chi-meraX.

## Acknowledgments

We thank members of the Carter and Niakan groups for their advice and support. We thank T. H. D. Nguyen, L. Passmore, D. Barford, T. Bharat and S. Chaaban for critical reading of the manuscript. We also thank S. Scheres, B. Toader and A.v. Kügelgen for advice on subtomogram averaging. We are grateful to the LMB Electron Microscopy Facility for access to, and support with, electron microscopy sample preparation and data collection. We thank J. Grimmett, T. Darling, and I. Clayson of LMB Scientific Computing for providing resources and A. Bukkosi from the LMB mechanical workshop for providing custom equipment. We thank F. Holfelder and T. Kohler for access to a confocal microscope, M. Shahbazi for access to a stereo microscope with a heated stage and E. R. Sanchez for assistance with mouse work. This work was funded by Medical Research Council as part of United Kingdom Research and Innovation MC_UP_A025_1011 (A.P.C), a European Molecular Biology Organization Postdoctoral Fellowship ALTF 426-2023 (K.H.) and a Wellcome Trust grant 221856/Z/20/Z (K.K.N.).

## Author contributions

K.S. and K.H. conceived the project, generated and analyzed the data. K.S., K.H. and A.C. wrote the initial draft. A.P.C., K.H. and K.K.N acquired funding. All authors reviewed and edited the manuscript.

## Data and materials availability

Atomic coordinates and cryo-EM maps have been deposited in the Protein Data Bank (PDB) or Electron Microscopy Data Bank (EMDB), respectively, under accession codes 9T63 and 55603 (Asymmetric unit of a CPL repeat), 9T64 and 55604 (CPL repeat unit with the central FBXW-SKP1 complex), 9T67 and 55607 (CPL repeat unit with ubiquitin bound), 9T66 and 55606 (Complex linking two CPL filament repeat units), 9T65 and 55605 (Complex linking two CPL filaments), 9T68 and 55609 (Composite structure of a CPL repeat unit) and EMDB-55608 (Subtomogram averaging map of CPL repeat unit structure from intact mouse embryos).

## Extended Data Figures and Tables

**Figure S1:**
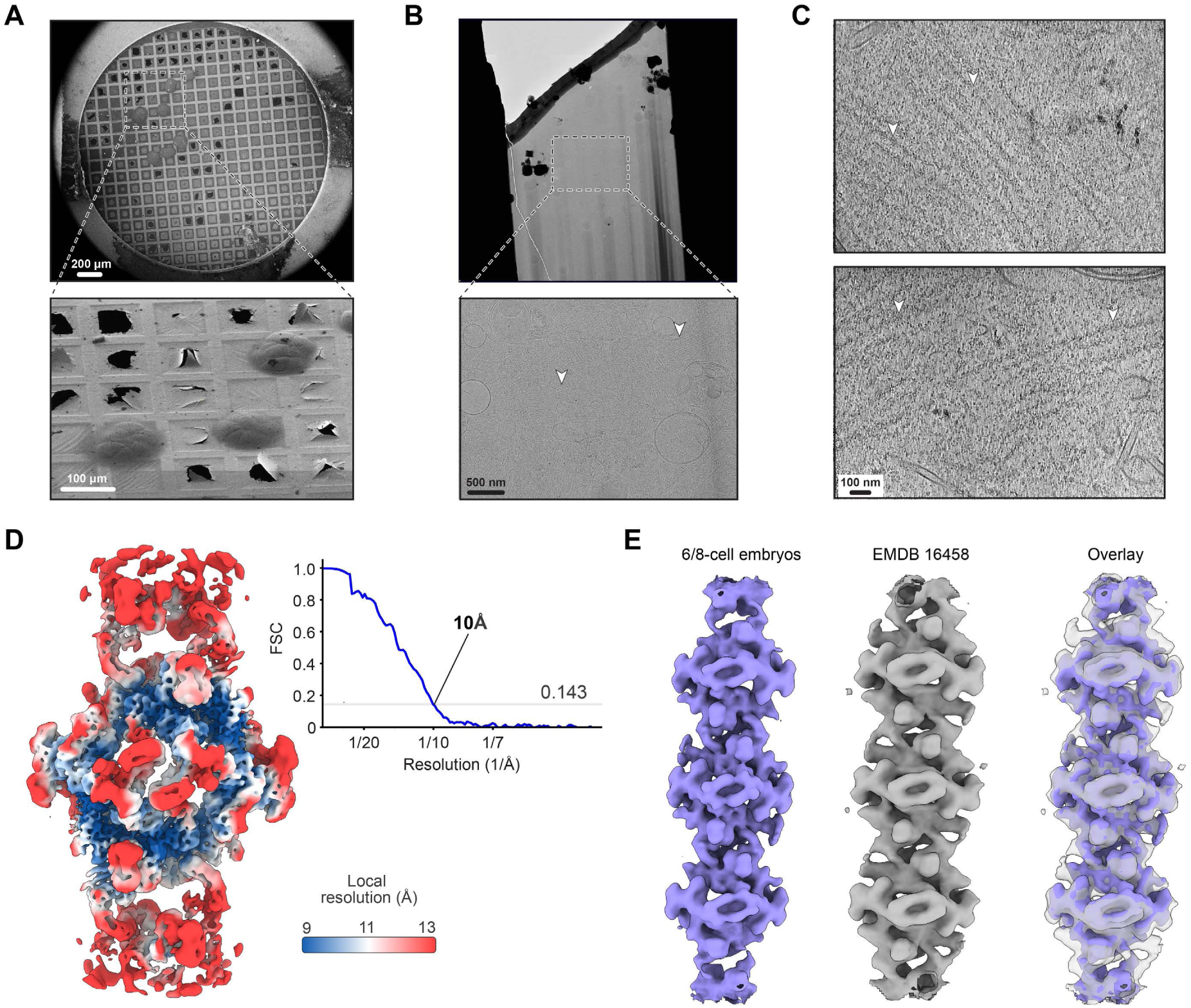
Structure of CPL filaments in 6/8-cell mouse embryos. **(A)** Scanning electron microscope (SEM) image of vitrified 6/8-cell mouse embryos on EM grids (top), and a FIB image showing three embryos on the grid (bottom). **(B)** Low-magnification transmission electron microscope (TEM) image of a representative lamella (top) and a zoomed-in view highlighting few examples of CPLs with white arrowheads (bottom). **(C)** Representative denoised tomographic slices showing CPLs with few examples highlighted with white arrowheads. **(D)** Subtomogram averaging map of the CPL repeat unit from cryo-ET data acquired on lamella of intact mouse embryos, coloured by local resolution. Corresponding Fourier shell correlation (FSC) curve is shown. **(E)** Comparison of CPL filament structure obtained using subtomogram averaging in early mouse embryos (left; this study) and from oocytes (center; EMDB-16458^27^). The two maps are overlaid (right), with EMDB-16458 rendered semi-transparent.

**Figure S2:**
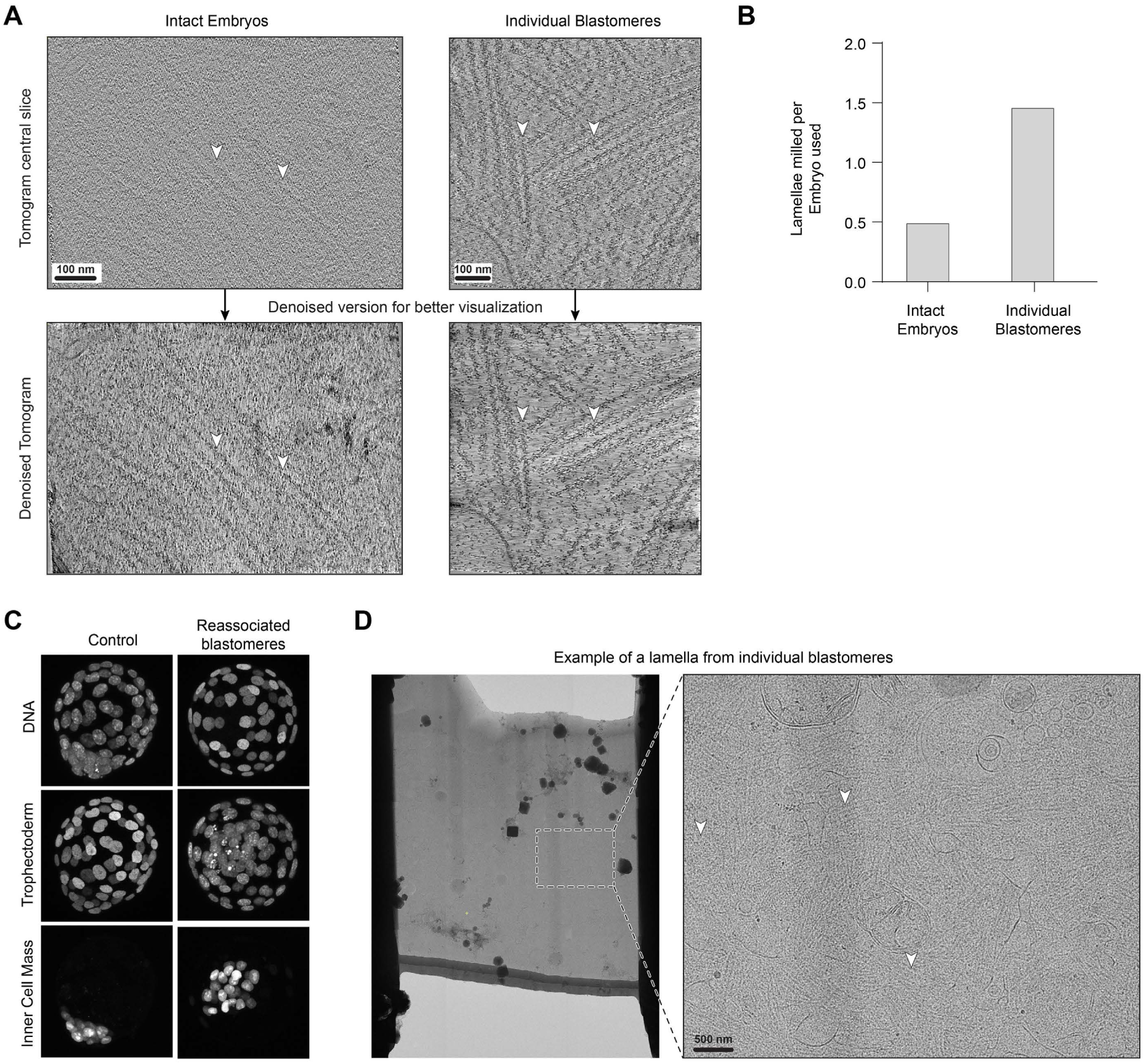
Cryo-ET of 6/8-cell embryo blastomeres. **(A)** Central slices of tomograms without and with denoising showing the improvement in contrast in data collected from individual blastomeres (right) as compared to that from intact embryos (left). Both tomograms were collected at a defocus of 4 µm and lamella thickness of 130-140 nm. **(B)** Graphs comparing the total number of lamellae obtained relative to the number of embryos used for vitrification on EM grids, for intact embryos (left) and embryos separated into individual blastomeres (right). Not all vitrified embryos/blastomeres yielded lamellae, as a fraction were lost during vitrification or located in regions of the grid unsuitable for cryo-FIB milling. **(C)** Immunofluorescence images showing the individual channels corresponding to the merged image in Figure 1D. **(D)** Low-magnification TEM image of a representative lamella prepared from a blastomere (left) and a zoomed-in view highlighting few examples of CPLs with white arrowheads (right).

**Figure S3:**
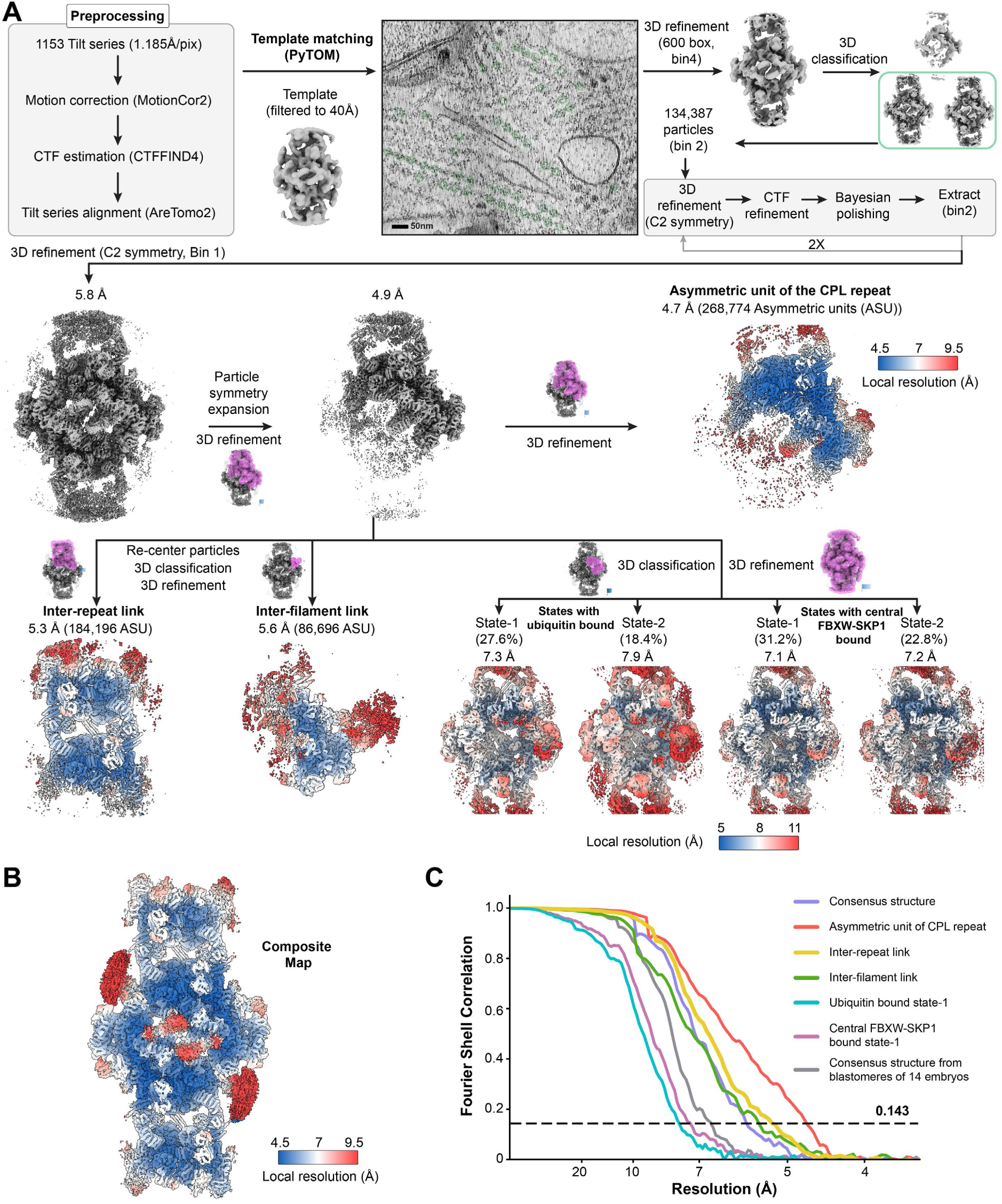
Subtomogram averaging processing pipeline. **(A)** The workflow for resolving the structure of the CPL repeat is shown. **(B)** Composite map generated by combining maps obtained after focussed 3D refinements is depicted and coloured based on local resolution. **(C)** Plot shows the gold standard Fourier Shell Correlation of the different subtomogram averaging maps shown in (A). The dotted horizontal line shows the 0.143 cut-off.

**Figure S4:**
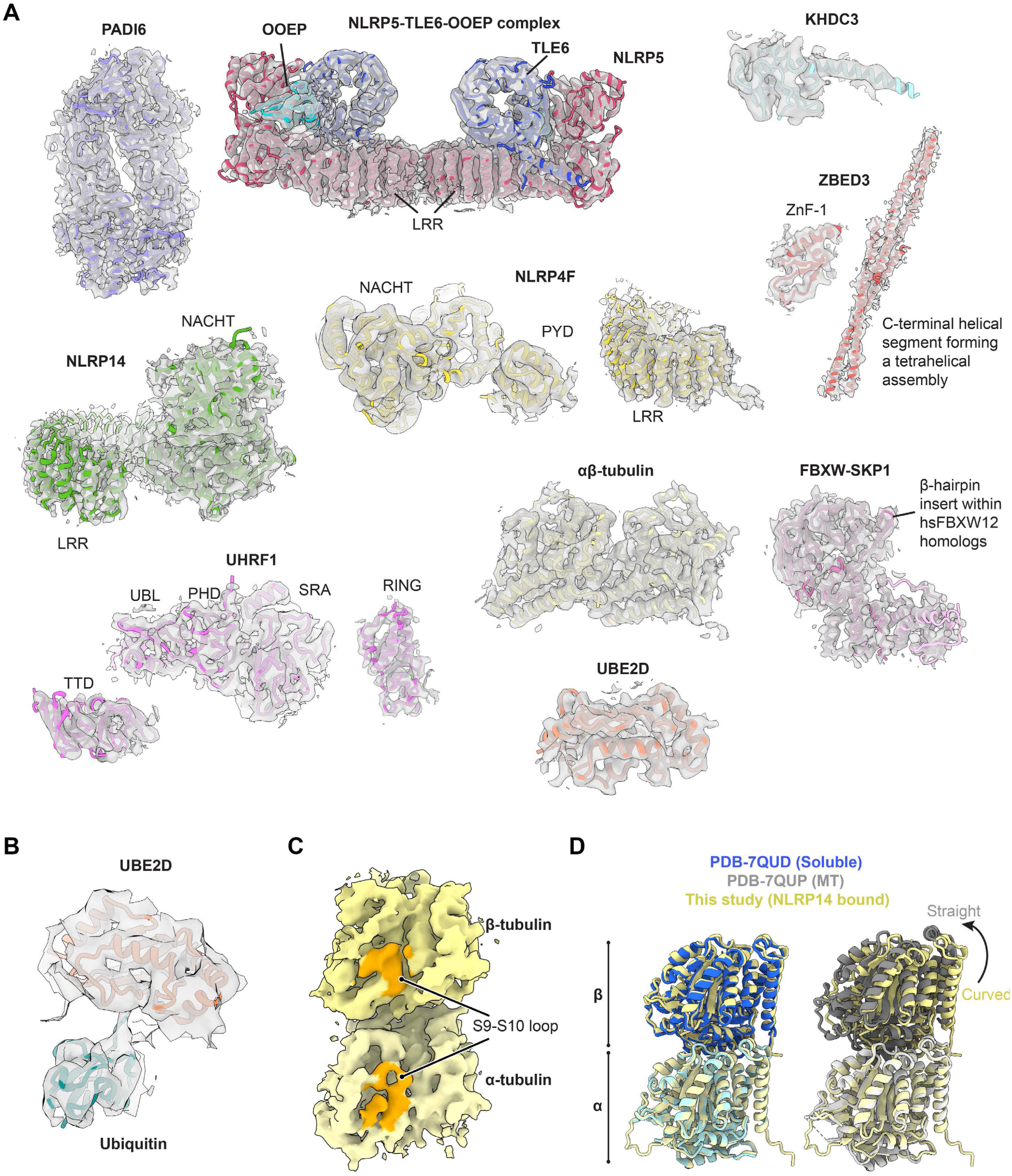
Densities of proteins identified within CPLs. **(A)** Gallery showing individual proteins with their atomic models fitted into the corresponding subtomogram averaging density maps. **(B)** Density corresponding to ubiquitin adjacent to UBE2D. **(C)** Density of the αβ-tubulin showing the S9-S10 loop in orange. The loop is shorter in the density assigned to β-tubulin as expected. **(D)** Structural comparison of αβ-tubulin from the CPL repeat with soluble (PDB 7QUD^78^) and microtubule-incorporated tubulin dimers (PDB 7QUP^78^), showing that CPL-bound αβ-tubulin adopts a conformation resembling the soluble form.

**Figure S5:**
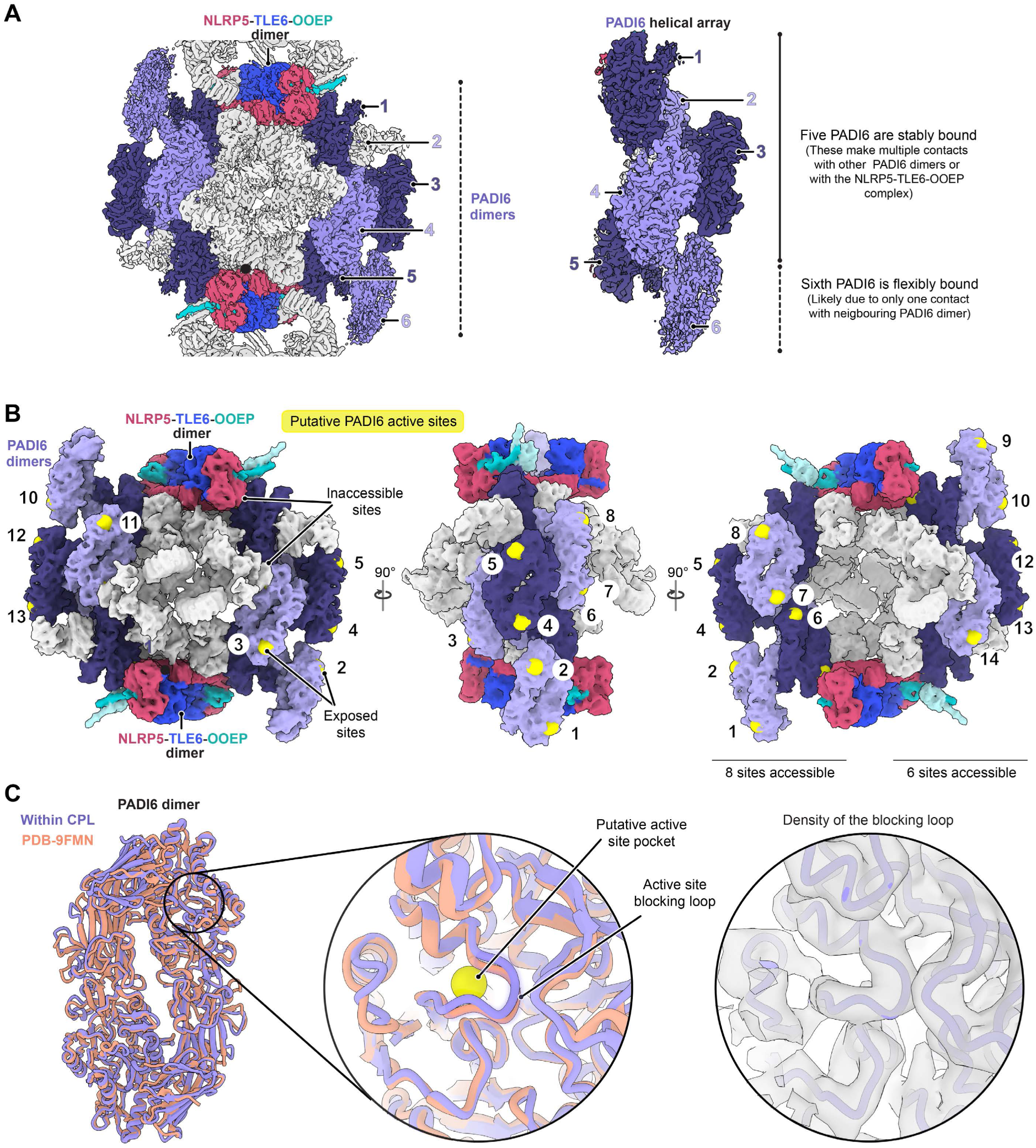
Arrangement of putative catalytic sites of PADI6 within CPLs. **(A)** Subtomogram averaging density of the PADI6 helical array is shown. Five PADI6 dimers (1-5) are stably bound whereas the sixth PADI6 dimer is more flexibly attached as its corresponding density is resolved at a lower resolution. **(B)** Putative catalytic sites of PADI6 within a CPL repeat unit are depicted as yellow spheres. A total of 14 sites are exposed whereas 10 sites are occluded by the interactions of PADI6 with the NLRP5-TLE6-OOEP complex or with the proteins bound to the central scaffold (shown in light grey). **(C)** The PADI6 dimer within the CPL helical array is compared with the structure of the isolated PADI6 dimer (PDB-9FMN)^44^. In both cases, the putative catalytic pocket is occluded. The subtomogram averaging density corresponding to the blocking loop is shown on the right.

**Figure S6:**
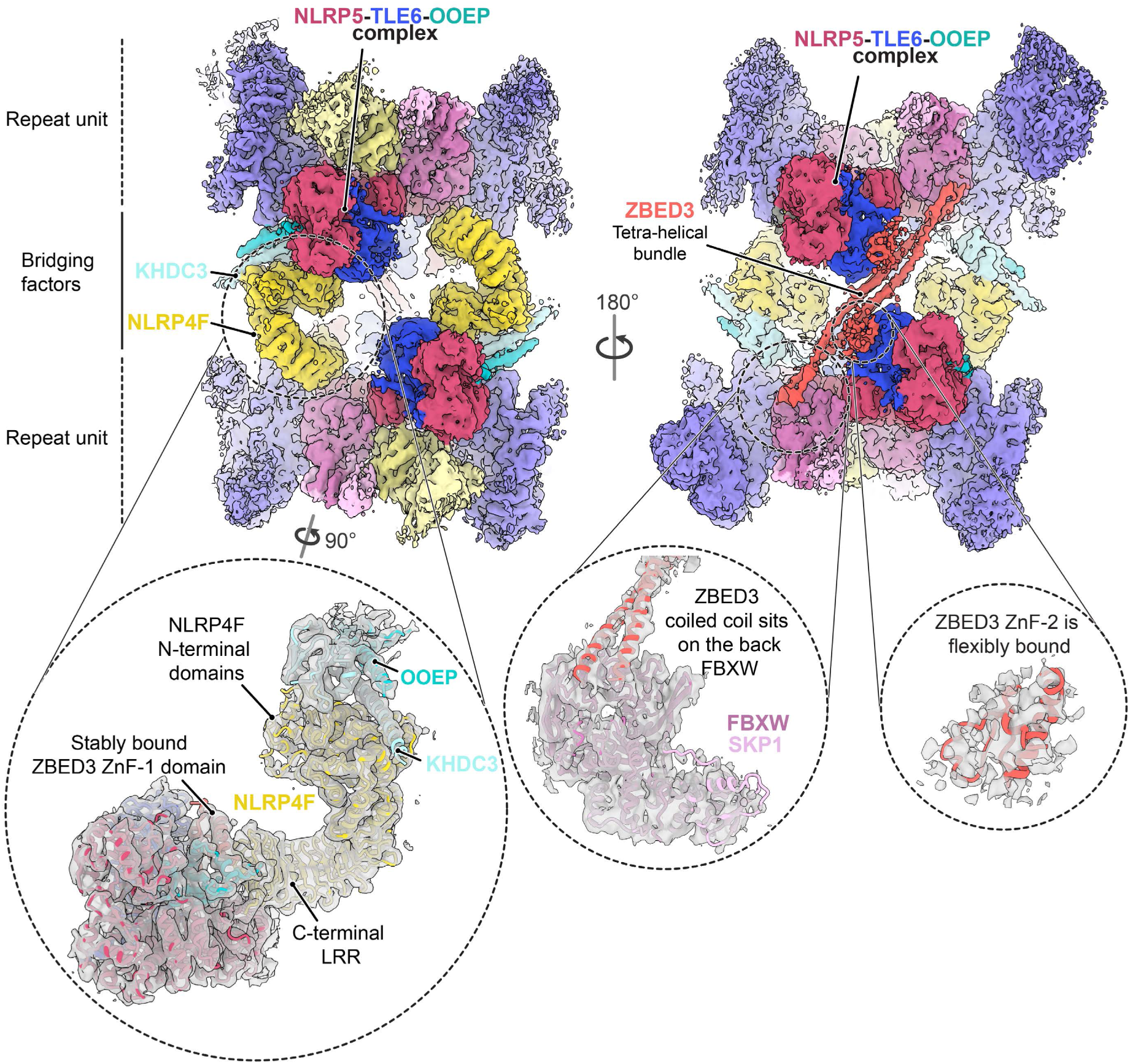
Subtomogram averaging density linking two CPL repeat units. Proteins mediating inter-repeat connections are labelled. Insets highlight NLRP4F, the interaction of the ZBED3 coiled-coil with a corner FBXW-SKP1 complex on the back face of the filament, and density corresponding to the flexibly attached ZBED3 zinc-finger (ZnF-2) domain bound to TLE6.

**Figure S7:**
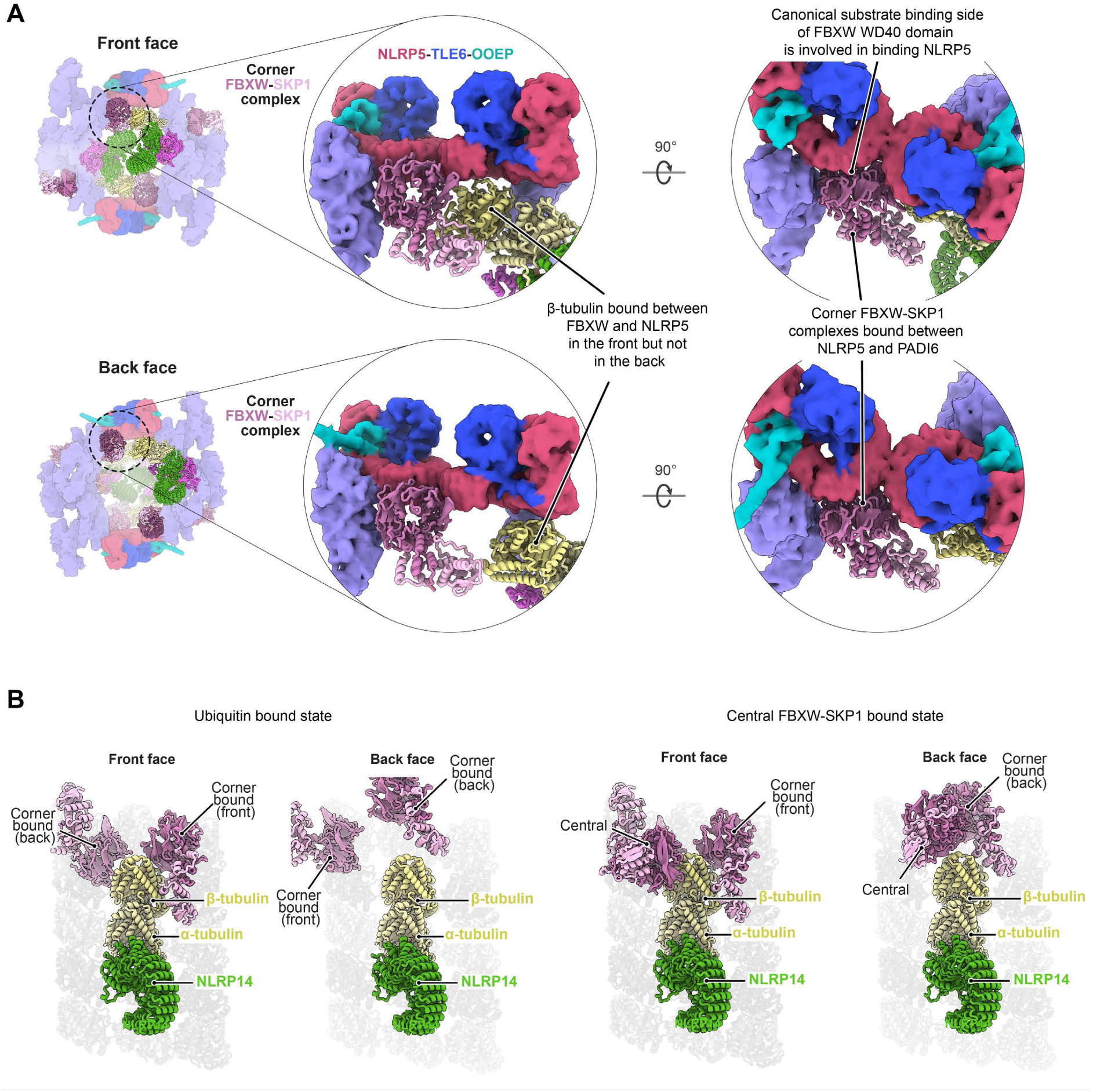
Interaction of FBXW-SKP1 complexes with UHRF1-UBE2D modules within the CPL repeat. **(A)** Models showing the interaction of corner FBXW-SKP1 complexes with the UHRF1-UBE2D modules on the front and back faces of the CPL repeat. On the front face, β-tubulin is positioned between FBXW-SKP1 and NLRP5, forming a compact interface that stabilizes the interaction with the UHRF1-UBE2D module. In contrast, on the back face the interface is more open and β-tubulin does not contact either component. Notably, the WD40 surface of FBXW-SKP1 that mediates substrate recognition in other F-box proteins is occluded through its interaction with NLRP5. **(B)** Interactions between αβ-tubulin-NLRP14 complexes on the front and back faces of the CPL repeat unit and FBXW-SKP1 complexes. The ubiquitin-bound state is shown on the left, and the state containing the central FBXW-SKP1 complex is shown on the right. To place these interactions in structural context of a microtubule, the tubulin heterodimer is overlaid onto a microtubule lattice.

**Figure S8:**
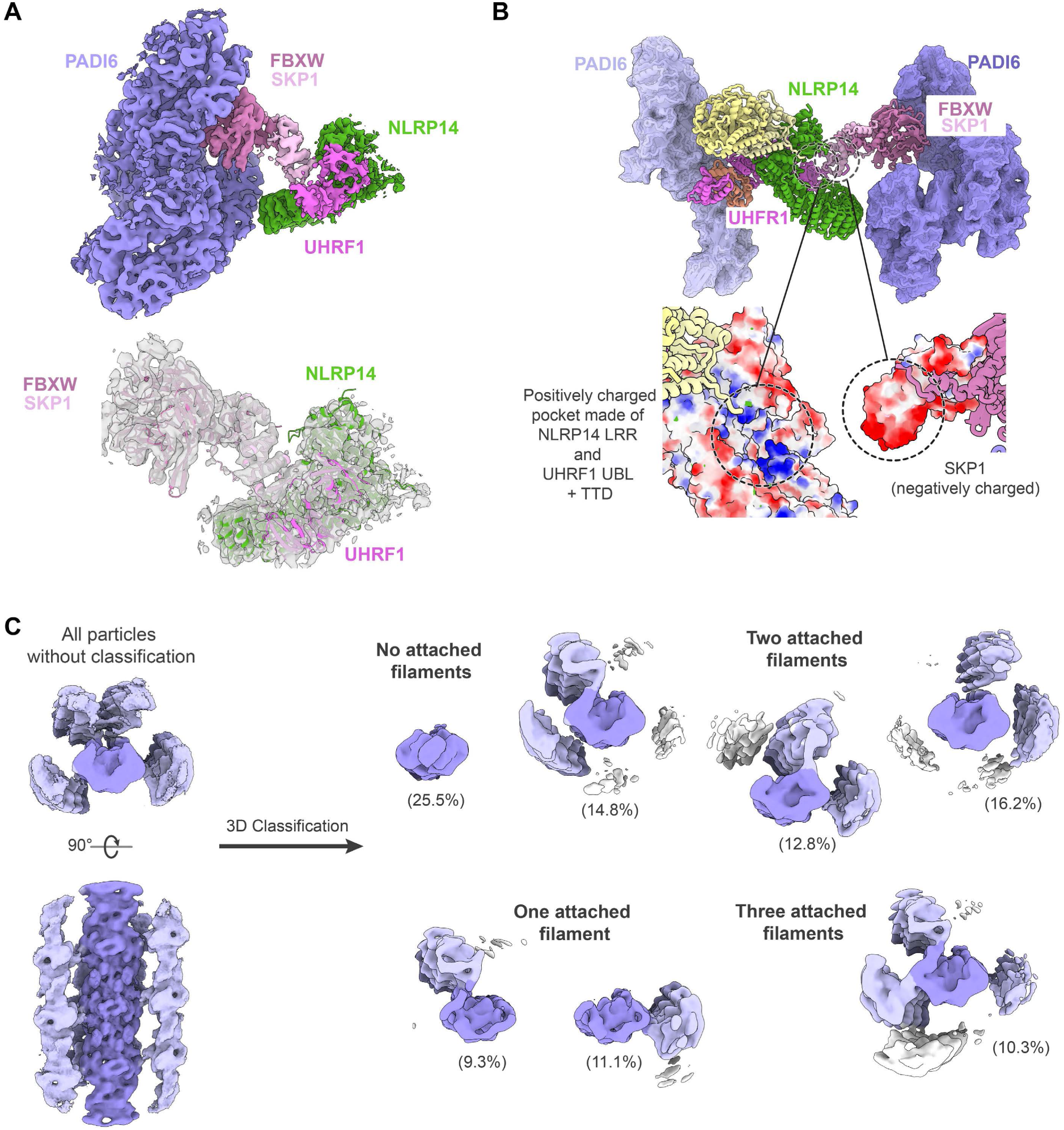
Inter-filament contacts within CPLs. **(A)** Subtomogram averaging density of the proteins linking two CPL filaments (top) with the corresponding atomic model of FBXW-SKP1, NLRP14, and UHRF1 fitted into the density (bottom). **(B)** View rotated 180° relative to Figure 6A showing that SKP1 engages the neighbouring CPL filament through a negatively charged patch (red) that binds a positively charged pocket (blue) formed by the NLRP14 LRR and the UHRF1 UBL and TTD domains. White colour denotes neutral charge. **(C)** Reconstruction of all particles corresponding to CPL repeats at a box size of 1,528 Å prior to 3D classification (left) and the resulting 3D classes after classification, with the percentage of particles assigned to each class indicated in brackets.

**Figure S9:**
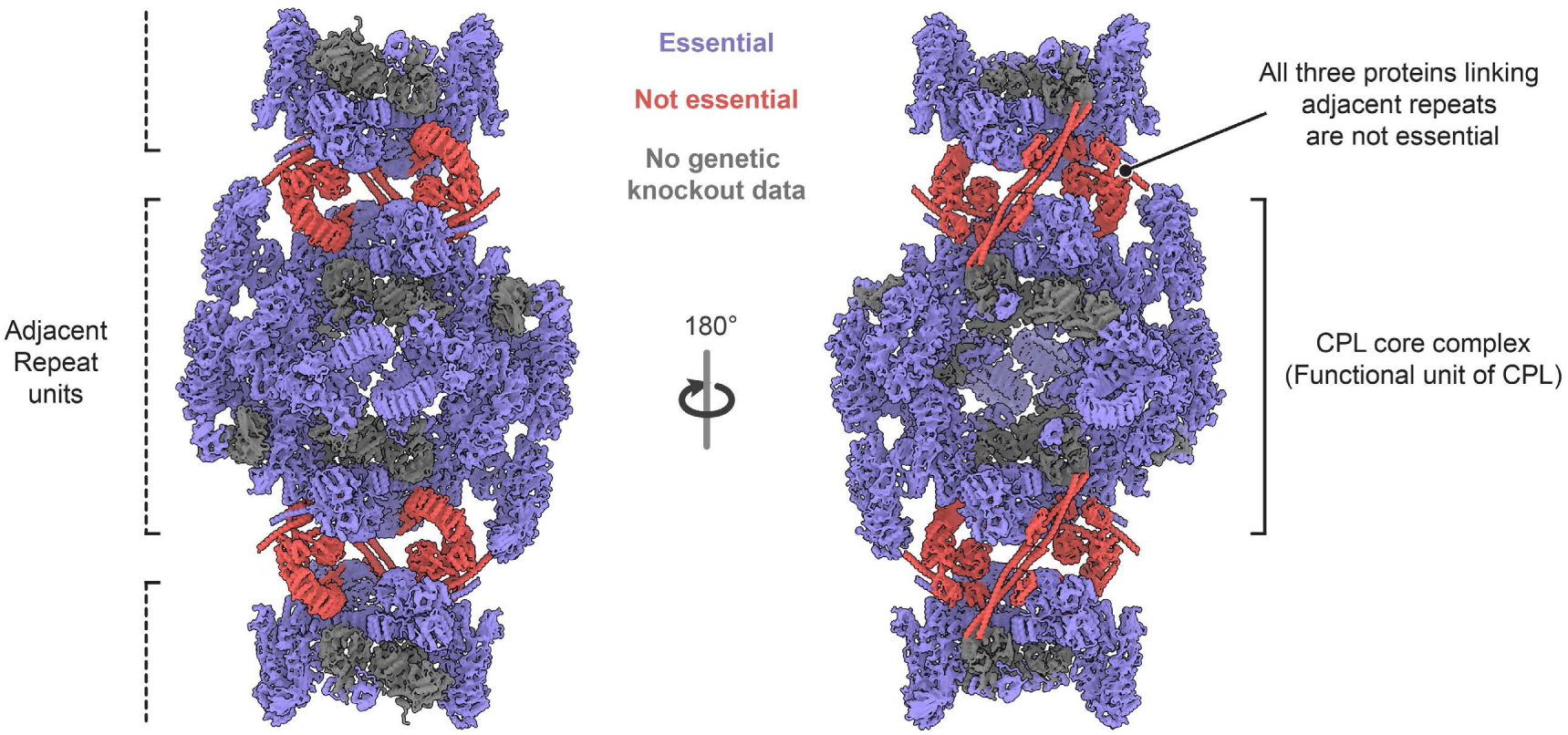
Mapping of genetic loss-of-function phenotypes onto the CPL structure. Proteins identified in the CPL repeat are coloured according to the developmental phenotypes reported upon gene disruption in mice. Proteins whose loss results in embryonic lethality or infertility are shown in lavender, proteins whose deletion permits viable progeny are shown in orange, and proteins for which genetic data are currently unavailable are shown in grey. This mapping highlights that factors essential for development form the CPL scaffold or reside within the CPL core complex, whereas proteins required for higher-order filament assembly are dispensable for viability.

**Figure S10:**
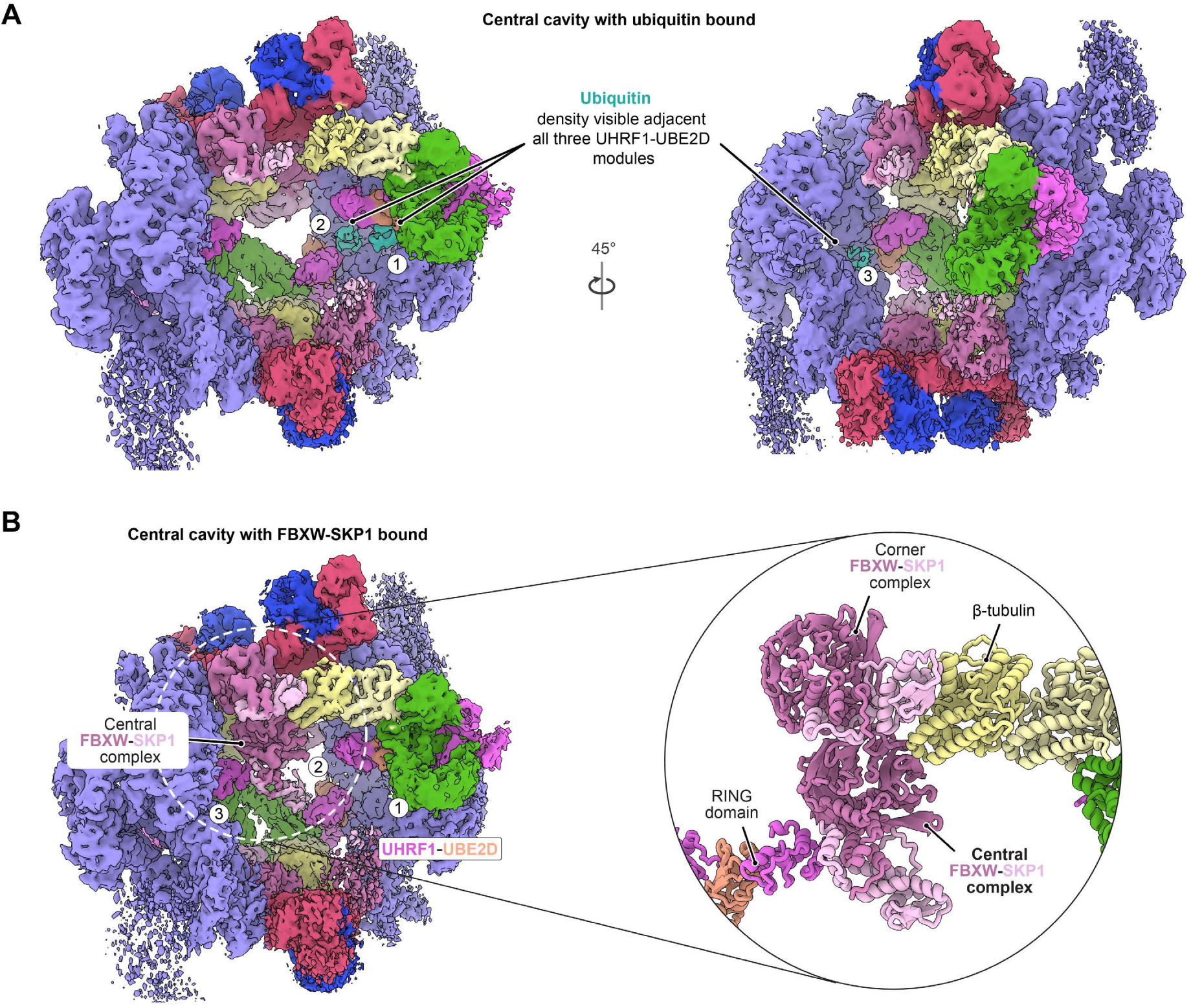
Two structural states of the CPL repeat unit. **(A)** Subtomogram averaging density of the CPL repeat unit showing density corresponding to ubiquitin adjacent to all three UHRF1-UBE2D modules within the central cavity, consistent with ubiquitin conjugated to UBE2D. **(B)** Density of the CPL repeat unit in a second state containing an additional central FBXW-SKP1 complex but lacking detectable ubiquitin density. In this state, the UHRF1-UBE2D module on the back face of the repeat shifts toward the central FBXW-SKP1 complex and contacts it via the β-tubulin.

**Figure S11:**
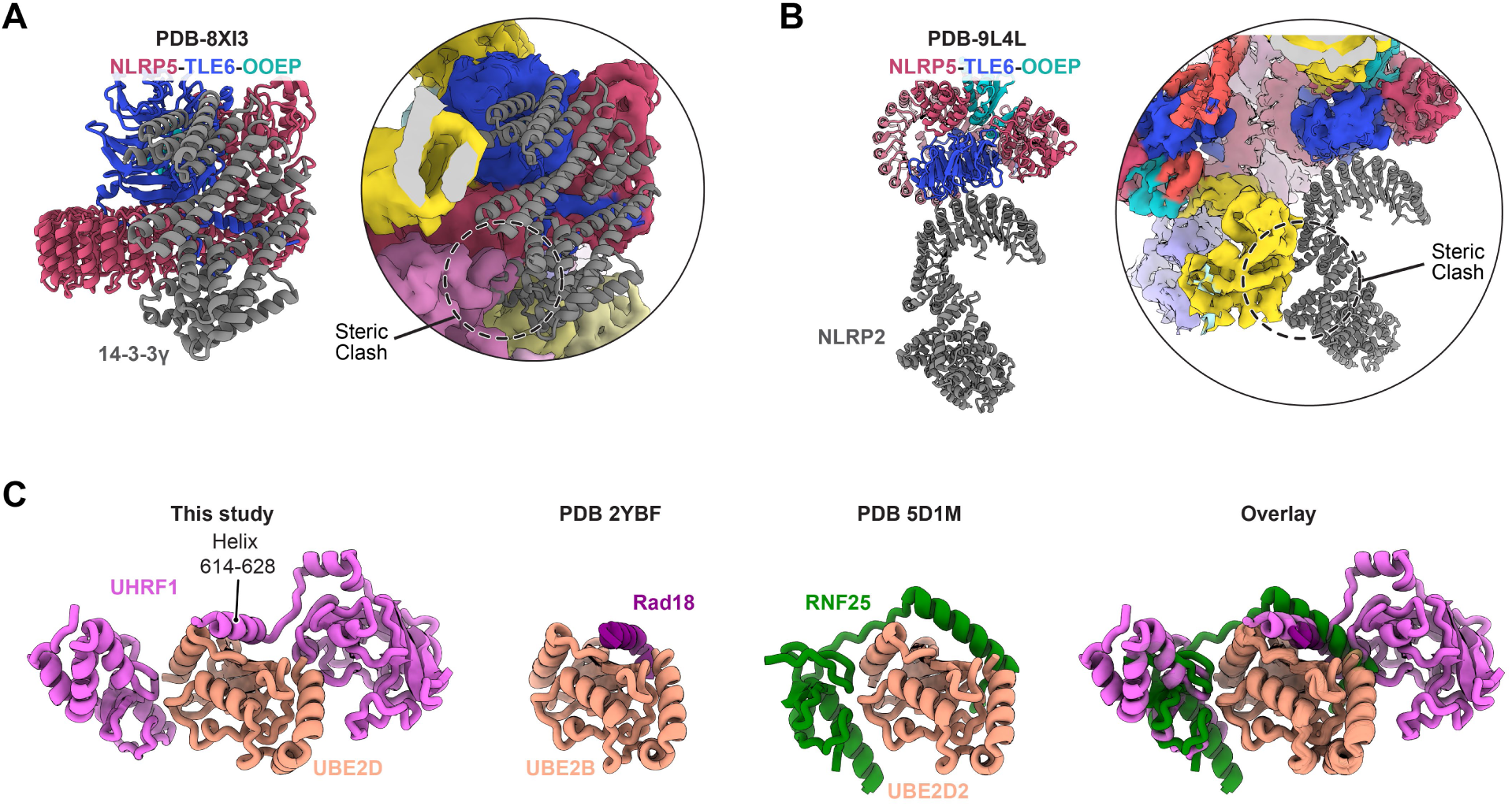
Comparisons with other structures. **(A-B)** Structural comparison of previously reported complexes containing the NLRP5-TLE6-OOEP complex bound to 14-3-3 protein or NLRP2 with the CPL structure. Superposition shows that incorporation of these factors into the CPL repeat would result in steric clashes, indicating that these interactions are incompatible with the assembled CPL architecture. **(C)** Comparison of backside interactions between E2 and E3 ligases. In the CPL structure, UHRF1 residues 614–628 form a helix that engages the backside surface of UBE2D. Similar helical backside interactions are observed in other E2–E3 complexes^58,59^ and are associated with promoting monoubiquitination or short ubiquitin chain formation.

**Table S1:** Cryo-ET data collection and model refinement statistics.

| Sample | Asymmetric unit of a CPL repeat | CPL repeat unit with central FBXW-SKP1 | CPL repeat unit with ubiquitin | Inter-repeat-link | Inter-filament-link | Composite model of a CPL repeat unit | CPL repeat unit from intact embryos |
| --- | --- | --- | --- | --- | --- | --- | --- |
| EMDB ID | 55603 | 55604 | 55607 | 55606 | 55605 | 55609 | 55608 |
| PDB ID | 9T63 | 9T64 | 9T67 | 9T66 | 9T65 | 9T68 | - |
| **Data collection and processing** |  |  |  |  |  |  |  |
| No. of Tomograms | 1,153 |  |  |  |  |  | 36 |
| Magnification | 105000X |  |  |  |  |  | 53000x |
| Voltage | 300 |  |  |  |  |  | 300 |
| Pixel size (Å) | 1.185 |  |  |  |  |  | 1.63 |
| Exposure (e<sup>-</sup>/Å<sup>2</sup>) per tilt (total tilts) | 3.5 (33) |  |  |  |  |  | 3.5 (33) |
| Frames collected | 8 |  |  |  |  |  | 6 |
| Defocus range (µm) | 2-4.5 |  |  |  |  |  | 2.5-4.5 |
| Final particle no. | 268774 | 34451 | 30490 | 184196 | 86696 | - | 7595 |
| Symmetry imposed | C1 | C1 | C1 | C1 | C1 | C1 | C1 |
| Map Resolution (Å) | 4.7 | 7.1 | 7.3 | 5.3 | 5.6 | - | 10 |
| FSC threshold | 0.143 | 0.143 | 0.143 | 0.143 | 0.143 | - | 0.143 |
| Sharpening B factor (Å<sup>2</sup>) | -180 | -200 | -200 | -150 | -100 | - | -200 |
| **Model Refinement** |  |  |  |  |  |  |  |
| Model Resolution (Å) | 6.0 | 7.7 | 8.0 | 6.3 | 6.5 | - | - |
| FSC threshold | 0.5 | 0.5 | 0.5 | 0.5 | 0.5 | - | - |
| **Model composition** |  |  |  |  |  |  |  |
| Non-hydrogen atoms | 76879 | 183153 | 181130 | 89612 | 35990 | 270155 | - |
| Protein residues | 15532 | 37008 | 36601 | 18104 | 7271 | 54588 | - |
| **Mean B factors (Å<sup>2</sup>)** |  |  |  |  |  |  |  |
| Protein | 206 | 317 | 352 | 264 | 257 | 269 | - |
| **RMSD deviations** |  |  |  |  |  |  |  |
| Bond lengths (Å) | 0.004 | 0.004 | 0.004 | 0.004 | 0.005 | 0.004 | - |
| Bond angles (°) | 1.12 | 1.109 | 1.099 | 1.119 | 1.121 | 1.118 | - |
| **Validation** |  |  |  |  |  |  |  |
| Clashscore | 1.06 | 1.97 | 2.12 | 1.54 | 2.16 | 1.92 | - |
| MolProbity score | 1.25 | 1.35 | 1.41 | 1.37 | 1.45 | 1.40 | - |
| **Ramachandran plot** |  |  |  |  |  |  |  |
| Favored (%) | 93.24 | 94.32 | 93.41 | 92.27 | 92.78 | 93.01 | - |
| Allowed (%) | 6.68 | 5.64 | 6.47 | 7.63 | 7.15 | 6.89 | - |
| Outliers (%) | 0.08 | 0.03 | 0.12 | 0.1 | 0.07 | 0.09 | - |

**Table S2:** Rationale for the placement of proteins within the CPL structure.

| Protein | Uniprot ID | Length | Molecular weight (kDa) | Modelled residues (copies per repeat) | Starting model | Rational |
| --- | --- | --- | --- | --- | --- | --- |
| PADI6 | Q8K3V4 | 682 | 76.78 | 1-682 (24) | AlphaFold3 prediction of PADI6 dimer | PADI6 was shown to co-localize with cytoplasmic lattices using immunofluorescence super-resolution microscopy<sup>27</sup> and is required for their formation<sup>79</sup>. Human PADI6 is a dimer<sup>44</sup>, hence we used AlphaFold3 to generate a model of the mouse PADI6 dimer. This model showed high correlation with the assigned density (Figure S4). |
| TLE6 | Q9WVB3 | 581 | 65.116 | 146-177, 247-581 (4) | PDB-8H93 | Shown to interact with each other and can form a dimeric complex<sup>7,30</sup>. TLE6 shown to co-localize with cytoplasmic lattices using immunofluorescence super-resolution microscopy<sup>27</sup> and all components are required for CPL formation<sup>7,9,10</sup>. The components show high correlation with the density. |
| OOEP | Q9CWE6 | 164 | 18.46 | 26-144 (4) | PDB-8H93; AlphaFold3 prediction was used to model the helical region after the KH domain. |  |
| NLRP5 | Q9R1M5 | 1163 | 131.32 | 97-470,485-1059 (4) | PDB-8H93 |  |
| KHDC3 | Q9CWU5 | 440 | 47.99 | 1-136 (2) | AlphaFold3 | Shown to interact with NLRP5-TLE6-OOEP complex and knockout leads to absence of CPLs<sup>7,32</sup>. The N-terminal 1-136 residues explained the density adjacent to OOEP consistent with previous binding data<sup>30</sup>. |
| ZBED3 | Q9D0L1 | 228 | 25.6 | 33-95,111-190 (4) | AlphaFold3 prediction of ZBED3 tetramer | Forms a complex with TLE6-OOEP-NLRP5 and knockout of this gene leads to the absence of CPLs<sup>23,24</sup>. The zinc finger domain fits into the density adjacent to the TLE6 WD40 domain and OOEP KH domain consistent with a recent in vitro structure<sup>29</sup>. Another density on the opposite side of the TLE WD40 also matched with the Zinc finger domain of ZBED3 but was resolved at a lower resolution. We have assigned this density to ZBED3. Adjacent to this was a coiled coil density that starts as a four helical bundle. The length of the coiled coil matches that of the ZBED3 C-terminal helix and AlphaFold3 predicted the formation of a four helical bundle by two ZBED3 dimers, which fits well into the corresponding density. |
| NLRP4F | L7N1W9 | 937 | 107.87 | 1-90,93-937 (2) | AlphaFold3 | Interacts with TLE6-OOEP-NLRP5 and knockout of this gene leads to the absence of CPLs<sup>23,24</sup>. Out of the other highly expressed NLRP proteins in early development<sup>80,81</sup>, NLRP4F has matching LRR length to that of the density. By contrast, other NLRP proteins, such as NLRP4A, 4B and 9b, have longer LRRs than the density suggests, whereas NLRP2 has a very short LRR region. NLRP4F levels also reduce drastically in TLE6 knockout oocytes<sup>27</sup> consistent with it interacting with the NLRP5-TLE6-OOEP complex. |
| NLRP14 | Q6B966 | 993 | 113.39 | 28-993 (3) | AlphaFold3 | Important for formation of CPLs and shown to form a complex with UHRF1<sup>11</sup>. One of the most abundant |
|  |  |  |  |  |  | NLRP proteins in oocyte and early embryos<sup>80,81</sup>. Has the matching LRR length when compared to the density. |
| UHRF1 | Q8VDF2 | 782 | 88.3 | 1-76,123-305,309-372,416-629,666-782 (3) | AlphaFold3 | Important for formation of CPLs and shown to form a complex with NLRP14<sup>11</sup>. Its domains show high correlation with the density adjacent to NLRP14. |
| UBE2D3 | P61079 | 147 | 16.69 | 1-147 (3) | AlphaFold3 | The protein levels of UBE2D3 reduce the most in PADI6 knockout oocytes as compared to other proteins with a similar fold<sup>27</sup>. UBE2D3 is the most abundant protein of the UBE2D protein family in oocytes and early embryos<sup>80,81</sup>. It also shows high correlation with the density. |
| α-tubulin | P68373 (Tuba1c) | 449 | 49.9 | 1-443 (3) | AlphaFold3 | Identified based on protein fold and matches the conformation of soluble tubulin heterodimer (PDB-7QUD<sup>78</sup>). The most abundant isoforms in mouse oocyte proteomics data<sup>81</sup> were built in the model but given the similarities between tubulin isoforms, the structure could contain different isoforms than those assigned. |
| β-tubulin | P68372 (Tubb4b) | 445 | 49.8 | 1-440 (3) | AlphaFold3 |  |
| FBXW | - | ~466 | ~53.59 | 1-466 (6-7) | AlphaFold3 prediction of FBXW15-SKP1 complex | The orthologs of human FBXW12 have a characteristic 8-propeller WD40 domains. FBXW15 is the only ortholog that is abundant in proteomic data from oocytes and embryos<sup>80,81</sup> was used as a template for model building. The density could also be explained by other orthologs of human FBXW12 in mice and is thus left unassigned as our density wasn't sufficient to determine side chain level differences. |
| SKP1 | Q9WTX5 | 163 | 18.67 | 1-163 (6-7) | AlphaFold3 prediction of FBXW15-SKP1 complex | Identified based on protein fold and shown to co-localize with CPLs using immunofluorescence super-resolution microscopy<sup>27</sup>. |

